# Integrating Natural and Engineered Genetic Variation to Decode Regulatory Influence on Blood Traits

**DOI:** 10.1101/2024.08.05.606572

**Authors:** Manuel Tardaguila, Dominique Von Schiller, Michela Colombo, Ilaria Gori, Eve L. Coomber, Thomas Vanderstichele, Paola Benaglio, Chiara Chiereghin, Sebastian Gerety, Dragana Vuckovic, Arianna Landini, Giuditta Clerici, Aurora Casiraghi, Patrick Albers, Helen Ray-Jones, Katie L. Burnham, Alex Tokolyi, Elodie Persyn, Mikhail Spivakov, Vijay G. Sankaran, Klaudia Walter, Kousik Kundu, Nicola Pirastu, Michael Inouye, Dirk S. Paul, Emma E. Davenport, Pelin Sahlén, Stephen Watt, Nicole Soranzo

## Abstract

Understanding the function of genetic variants associated with human traits and diseases remains a significant challenge. Here we combined analyses based on natural genetic variation and genetic engineering to dissect the function of 94 non-coding variants associated with haematological traits. We describe 22 genetic variants impacting haematological variation through gene expression. Further, through in-depth functional analysis, we illustrate how a rare, non-coding variant near the *CUX1* transcription factor impacts megakaryopoiesis through modulation of the *CUX1* transcriptional cascade. Collectively, our findings enhance the functional interpretation of genetic association studies and advance understanding of how non-coding variants contribute to blood and immune system variation.

## Introduction

Genome-wide association studies (GWAS) of haematological traits have identified thousands of variant–phenotype associations that illuminate key aspects of blood homeostasis and immune function^1,2^. Large-scale quantitative trait locus (QTL) studies in immune cells^3^ and whole blood^4^ have been instrumental in explaining how GWAS-supported genes influence blood traits in health and disease^2,5^. Notably, up to 90% of GWAS associations for haematological phenotypes lie in non-coding regions of the genome^2^, complicating the identification of target genes and the elucidation of the mechanisms through which these variants act. Understanding the transcriptional regulators mediating such effects is crucial for therapeutic discovery, as exemplified by the recent approval of CRISPR/Cas9-based ex vivo therapies (Casgevy) for sickle cell disease and β-thalassemia^6^. In this context, the identification of non-coding variants controlling fetal haemoglobin levels through regulation of the transcriptional repressor BCL11A^7,8^ led to the discovery of an erythroid-specific enhancer that became a successful therapeutic target.

Variant-to-function studies remain challenging due to limitations in scaling functional validation, prioritising variants within extended linkage disequilibrium (LD) blocks, and establishing definitive variant–gene relationships. Functional annotation of haematological variants has relied primarily on expression QTL (eQTL) analyses^4,5^ and high-throughput experimental approaches such as Massively Parallel Reporter Assays (MPRA)^9–11^, CRISPR/Cas9^12^ and CRISPR interference (CRISPRi)^13^. Despite the insights gained from these methods, several challenges remain. First, the functional impact of the growing number of rare non-coding GWAS variants (RNVs) revealed by dense imputation and whole-genome sequencing (WGS)^14^ remains insufficiently explored. eQTL studies are underpowered for rare variants, and few MPRA^10^ and CRISPRi^13^ screens have included them in their designs. Second, discrepancies between *in vivo* expression studies and *in vitro* functional assays remain largely unresolved, as the latter often test variants outside their native chromatin context (e.g., episomal MPRAs^10^), potentially introducing experimental artefacts^15,16^. Additionally, phased WGS data, which are essential for dissecting haplotype-specific effects, are often lacking in eQTL studies. Finally, few functional screens have been conducted in cell types directly relevant to the blood phenotypes associated with these variants^17^.

Here, we address the regulatory function of 94 rare non-coding variants associated with haematological traits using an MPRA to assess enhancer activity, together with analyses of differential gene expression (DE) and alternative transcript usage (ATU) in 2,971 whole-blood RNA-seq samples with phased WGS data. Through extensive manual curation, we identify 22 variants with direct regulatory evidence in genes robustly associated with blood traits. Finally, we perform an in-depth functional analysis of two of these variants, elucidating a molecular mechanism that recapitulates its GWAS association.

## Results

### Variant prioritisation and MPRA in hematopoietic cell lineages

To identify candidate regulatory variants affecting haematological traits, we began with 12,181 loci associated with 29 blood phenotypes from a GWAS conducted by our group^2^. These traits (**Table S1**) encompass a wide range of clinical indices reflecting erythroid, megakaryocytic, monocytic, and lymphoid lineages. From the 178,890 variants contained within the 95% fine-mapping credible sets, we applied sequential filters. First, we restricted the dataset to rare variants

(minor allele frequency [MAF] ≤ 1% in the UK Biobank), retaining 5,813 variants, and then excluded coding variants using the most severe consequence annotation from the Variant Effect Predictor (VEP; **Methods**), yielding 5,248 non-coding variants. We next selected variants with at least one association fine-mapped at high posterior probability (PPFM ≥ 0.9), narrowing the set to 196 variants, and finally prioritised those with large effect sizes (β < first or > third quartile of the standardised trait distribution) (**Figure 1A, Supplementary Figure 1A–B, Methods)**. These criteria identified 94 rare non-coding variants (RNVs), hereafter referred to as “index variants” (**Figure 1A, Table S2)**.

**Figure 1.**
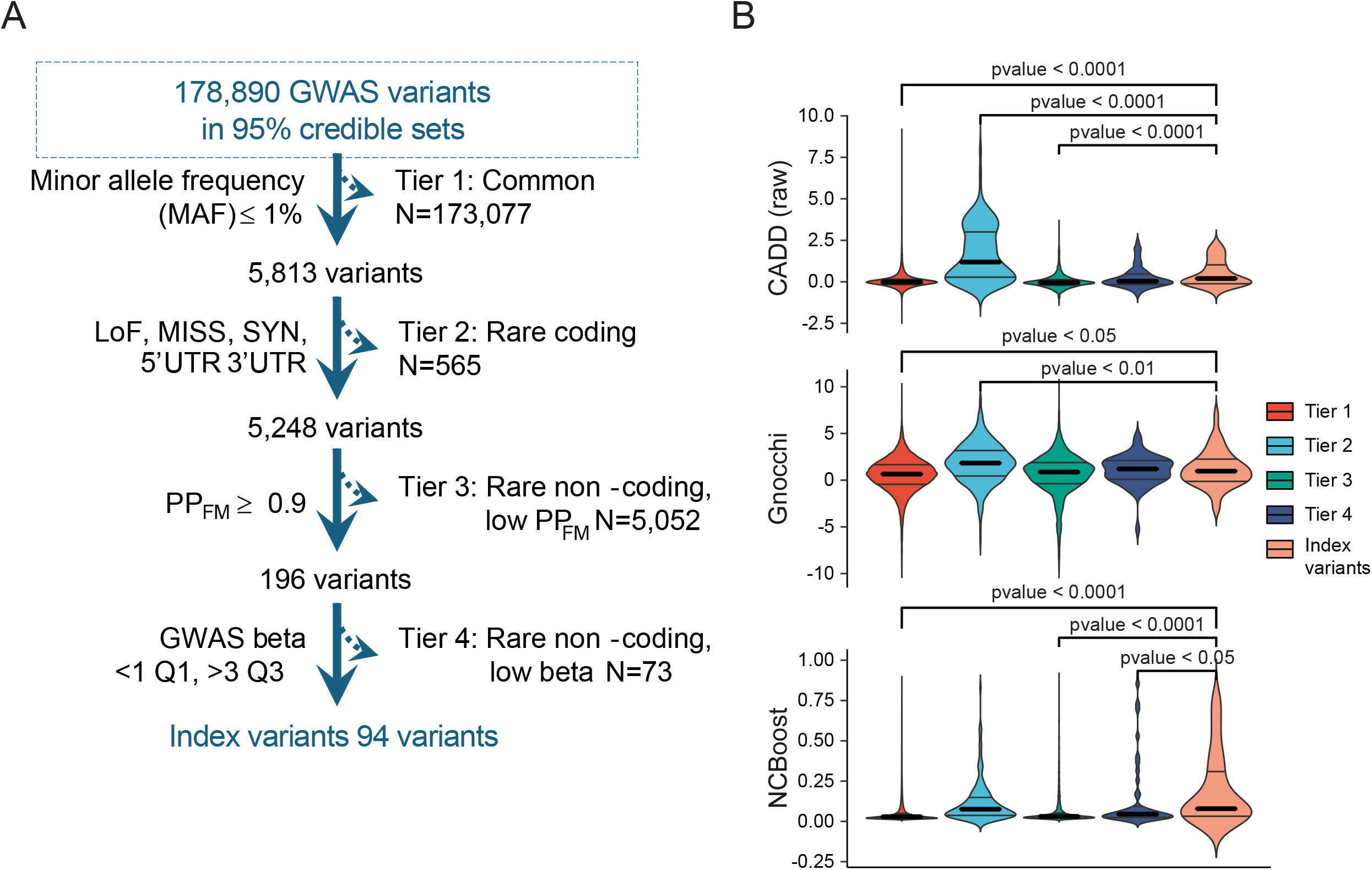
Variant prioritisation strategy and MPRA controls and inter-replica variability. **A)** Prioritisation strategy and **B)** values of the CADD, Gnocchi and NCBoost scores across the different tiers. *The comparisons correspond to the values of the ‘index variants’ versus the rest of the tiers, Wilcoxon test*.

The index variants showed significantly larger effect sizes than heterozygous pathogenic ClinVar/HGMD variants associated with blood traits^2^ (p-value = 0.007, *Wilcoxon test*). Notably, 95% of the index variants remained significantly associated with at least one blood trait after conditioning on common variants. Moreover, index variants displayed significantly higher values of orthogonal prioritisation scores—including Combined Annotation Dependent Depletion (CADD]^18^), NCBoost^19^, and genomic non-coding constraint of haploinsufficient variation ([Gnocchi]^20^) compared with other RNV classes (**Figure 1B)**.

We incorporated the 94 index variants into an MPRA library designed to test enhancer activity^15,21^ together with 15 positive and four negative controls selected from previous studies^9,22^. To explore different sequence contexts^23^, each allele was synthesised within five partially overlapping tiles, each tagged with unique barcodes, and cloned into an MPRA reporter vector (**Methods**). The resulting library of 19,050 oligonucleotides was transfected in seven replicates into four blood-derived cancer cell lines: K-562 (chronic myeloid leukaemia; erythroid model), CHRF-288-11 (acute megakaryoblastic leukaemia; megakaryocyte model), HL-60 (acute myeloid leukaemia; neutrophil model), and THP-1 (acute monocytic leukaemia; monocyte model) (**Figure 2A**).

**Figure 2.**
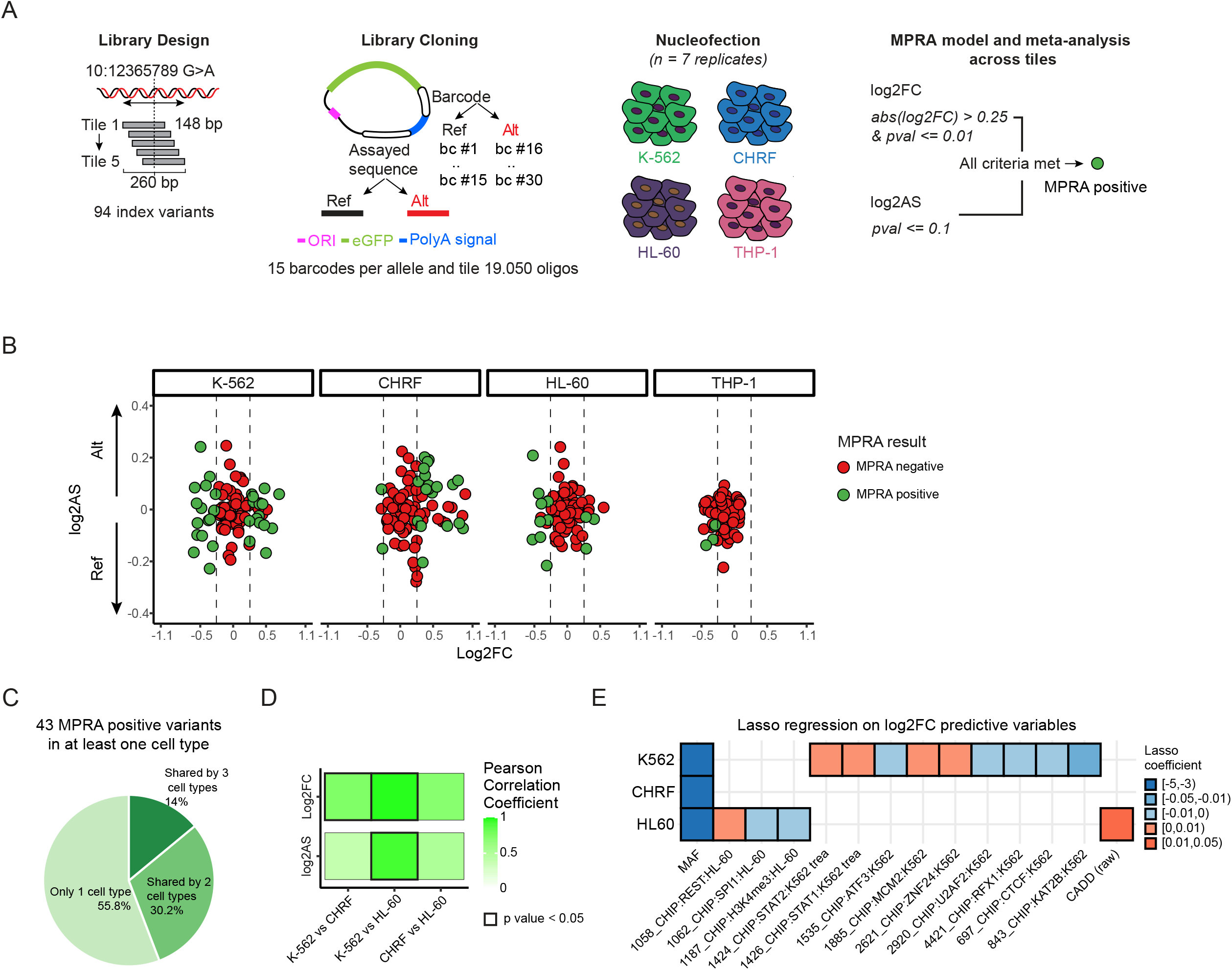
MPRA screening. **A)** Design of the MPRA for enhancer activity and **B)** results after the meta-analysis. **C)** Sharing of MPRA positive variants between the cell types assayed. **D)** Pearson correlation coefficients for MPRA positive variants across K-562, CHRF-288-11 and HL-60 cells. THP-1 had too few shared MPRA positives to perform meaningful comparisons. **E)** Lasso regression on sequence features predictive of Log2FC.

Enhancer activity and allele-specific expression were measured using *MPRAmodel*^24^ respectively from log_2_ fold change (log_2_FC) and log_2_ allelic skew (log_2_AS) measurements for each tile, variant, and cell type (**Table S3**). Meta-analysis across overlapping tiles produced summary log_2_FC and log_2_AS values per variant and cell type^10^ (**Table S4**). The directionality of allelic skew in positive controls showed strong concordance with prior MPRA data in K-562 cells^9^ (linear model R^2^ = 0.821, **Supplementary Figure 1C**). We identified 43 variants that significantly modulated enhancer activity, henceforth referred to as *MPRA-positive* (MPRA^+^, **Figure 2A–B, Supplementary Figure 1D, Tables S4 and S6**). Among the four cell lines, THP-1 yielded the fewest MPRA^+^ variants (n = 4; **Supplementary Figure 1E**), likely due to lower nucleofection efficiency^25^ as reflected by the proportion of GFP-positive cells post-transfection (median 55.5% in CHRF-288-11 vs. 6.8% in THP-1; *p* = 0.01, Wilcoxon test).

Of the 43 MPRA^+^ variants, 25 were specific to a single cell type, whereas fourteen and four were shared across two and three cell types, respectively (**Figure 2C, Supplementary Figure 1E**). We observed high correlation (Pearson *r* > 0.9) in log_2_FC and log_2_AS between MPRA^+^ variants shared by K-562 and HL-60 cells (**Figure 2D, Supplementary Figure 1F**), consistent with their shared myeloid leukaemia origin^26,27^.

To identify factors influencing enhancer activity, we integrated cell-matched sequence features from Enformer^28,23^ with GWAS-derived parameters and applied lasso regression^23^. Among variant features, allele frequency was a significant predictor of enhancer activity in K-562, CHRF-288-11, and HL-60 cells (**Figure 2E**), with rarer variants (MAF < 0.01) having higher average log_2_FC values compared to common variants (**Supplementary Figure 1G**), although additional data will be required to confirm this trend. Higher CADD scores were also predictive of elevated log_2_FC values in HL-60 cells (**Figure 2E**). Analysis of CHIP-seq data further showed that motifs bound by activating transcription factors (e.g., *STAT1, STAT2*) were associated with positive log_2_FC values, whereas those bound by repressors or insulators (e.g., *CTCF, RFX1*) correlated with negative values (**Figure 2E**).

Collectively, these findings demonstrate that index variants affect enhancer activity in our cell systems, and that MPRA scores reflect population characteristics of the genetic variants.

### Population survey of RNA expression in healthy volunteers supports variant effects mediated by transcription

To evaluate the regulatory impact of the index variants within their native chromatin context, we analyzed two bulk RNA-seq datasets (**Table S5**). The first dataset, from the INTERVAL study^29^, includes gene and transcript quantification from whole-blood samples of 2,971 individuals with WGS data (mean coverage = 15×). Overall, we identified heterozygous carriers for 88 of the 94 index variants, with a median of 40 carriers per variant (range = 3–163 individuals). Because the traditional eQTL approach^4^ is underpowered for variants with low allelic counts, we applied a calibrated differential gene expression (DE) analysis that accounted for multiple experimental covariates to test for expression differences between wild-type and heterozygous carriers of rare alleles (**Methods**). In parallel, we examined regulatory mechanisms beyond gene-level control by testing for alternative transcript usage (ATU), defined as changes in the relative abundance of expressed transcripts for each gene^30^. ATU modeling was performed using an additive log-ratio approach that accounts for compositional structure and incorporates all experimental covariates used in the DE analysis (**Methods**).

We detected significant DE or ATU effects for 42 of the 88 variants, involving 60 genes (**Figure 3A, Table S5**). Among these, 23 variants (29 genes) showed only DE effects, 11 variants (11 genes) exhibited only ATU effects, and 8 variants displayed both types of regulation (affecting 14 genes with DE, 4 with ATU, and 5 with both DE and ATU; **Table S5**). As additional evidence for transcript-level regulation among the ATU genes, 8 of 20 showed splicing QTL (sQTL) signals in whole blood from the GTEx Project^31^, although these involved common variants (MAF > 1%) that were conditionally independent of those tested here (r^2^ > 0.7, EUR population, 0.5 Mb window; **Methods**). Furthermore, we identified three ATU variants as sQTLs for their corresponding genes and observed common sQTLs for an additional 11 ATU genes in the INTERVAL QTL repository^29^. One notable example involved the synonymous cryptic splicing variant rs150813342 in *GFI1B*, a transcriptional repressor critical for platelet and red blood cell development (**Supplementary Figure 2A**). Editing this variant using CRISPR/Cas9 in K-562 cells produced comparable transcript-usage alterations^31^.

**Figure 3.**
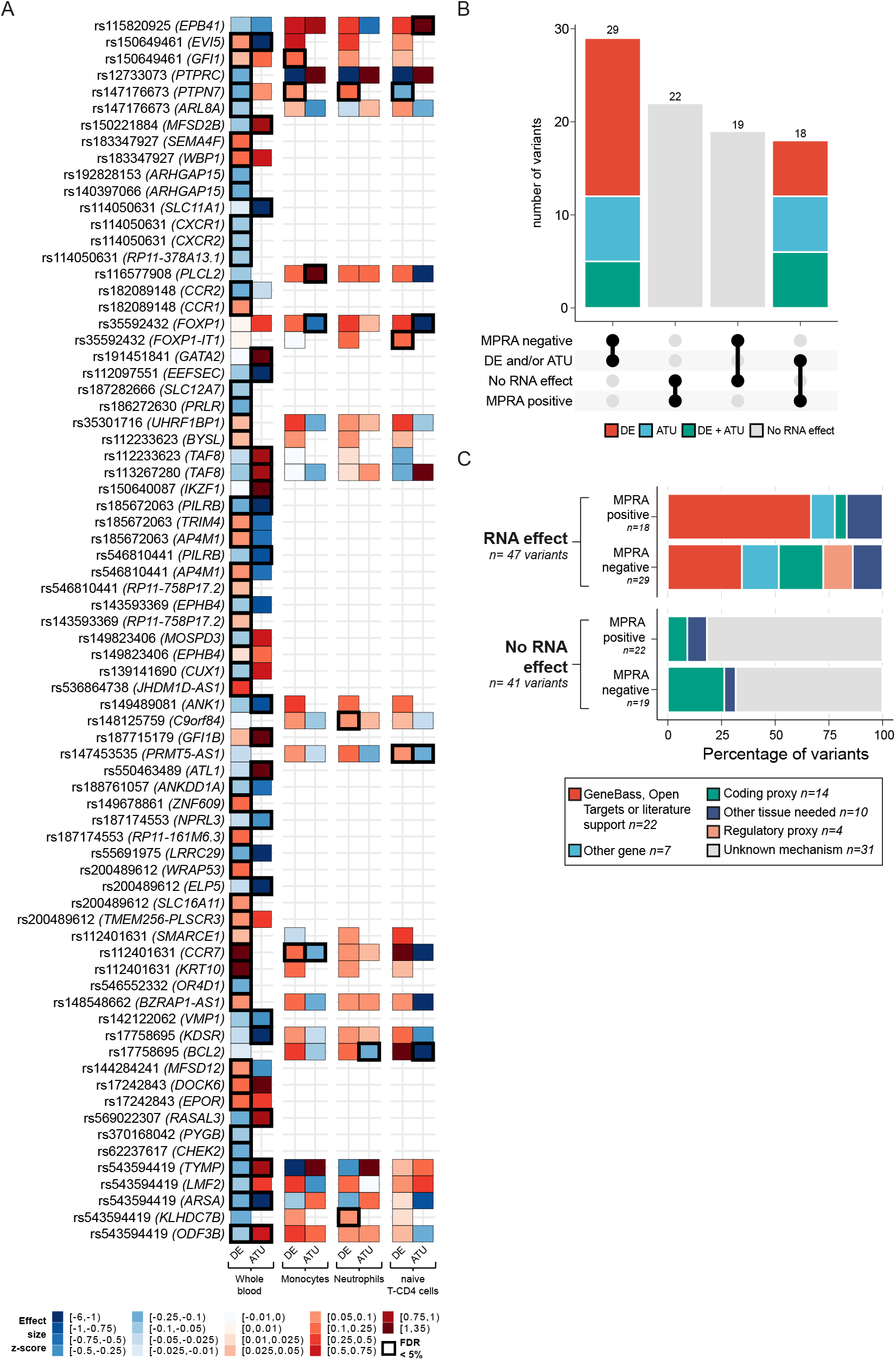
RNA-seq study results and curation of screened variants derived from RNA Seq and MPRA readouts. **A)** Variants with significant DE/ATU genes. **B)** Breakdown of the overlap between the results of the MPRA and the RNA-seq. **C)** Breakdown of the mechanistics and manual curation labels by the results of the MPRA and the RNA-seq study.

To dissect cell-type-specific contributions, we next analyzed DE and ATU in three immune cell types (monocytes, neutrophils, and naïve CD4^+^ T cells) from the BLUEPRINT human variation panel^3^. Of the 88 variants assessed in INTERVAL, 25 had at least one heterozygous carrier among the 196 BLUEPRINT donors (median = 5 carriers per variant; range = 1–17). We identified DE or ATU evidence for 10 of these 25 variants, involving 11 genes (**Figure 3A, Table S5**). Among these, four variants (four genes) showed only DE effects, three (three genes) showed only ATU effects, and 3 displayed both types of regulation (affecting one gene with DE, one with ATU, and two with both DE and ATU). Of the ten BLUEPRINT DE/ATU variants, five overlapped with the INTERVAL findings, three involving the same genes. In four of the remaining five cases, regulatory effects were restricted to a single BLUEPRINT cell type (**Figure 3A, Table S5**). Collectively, these RNA-seq data support multiple MPRA^+^ variants, highlighting both shared and context-specific regulatory mechanisms.

### Regulatory landscapes defined by *in vivo* and *in vitro* studies

We integrated evidence from the MPRA and RNA-seq (DE/ATU) experiments to determine how multiple lines of evidence converge toward a single mechanistic interpretation for each genetic association. Based on combined results, the 88 tested variants were categorized into four groups: 18 *double-positive* variants (regulatory effects in both MPRA and RNA-seq), 22 MPRA^+^/RNA^−^, 29 MPRA^−^/RNA^+^, and 19 MPRA^−^/RNA^−^ variants (**Figure 3B, Table S6**).

Within the double-positive set, we evaluated the concordance in the direction of the effect allele between MPRA allelic skew and DE and found moderate agreement (8/14 cases; **Methods**). Among the 22 MPRA^+^/RNA^−^ variants (**Figure 3B**), those lacking RNA-seq effects had significantly fewer heterozygous carriers than DE and/or ATU variants (p = 0.045, Wilcoxon test; **Supplementary Figure 2B**). We next assessed whether the 29 MPRA^−^/RNA^+^ variants were enriched for ATU cases that might evade detection in enhancer assays; however, no significant difference in ATU distribution was observed between double-positive and MPRA^−^/RNA^+^ variants (p = 1, χ^2^ test; **Figure 3B, Table S6**). To identify sequence features distinguishing MPRA^−^/RNA^+^ from double-positive variants, we applied lasso regression (**Methods**) and detected three ChIP-seq motifs in K-562 cells—*E2F, RLF*, and *BCLAF1*— associated with the MPRA^−^/RNA^+^ class (**Supplementary Figure 2C**). Notably, both *E2F* and *RLF* showed very low expression in K-562 cells (**Supplementary Figure 2D**).

To annotate potential effector genes underlying these associations, we integrated information from the GeneBass^32^ and OpenTargets^33^ databases and performed a detailed literature review (**Supplementary Figure 3, Table S6**). For variants with RNA-seq regulatory evidence, all DE/ATU genes were considered; for RNA-seq–negative variants, we included all tested genes. In total, genes were annotated for 76 of the 88 variants.

Using phased WGS data (unavailable at the time of the MPRA design), we identified cases where nearby coding variants in high linkage disequilibrium (LD) with the index variant (r^2^ = 0.26–0.97) could explain the observed association, labelling these as “coding proxies”. Fourteen variants had at least one such proxy, and regulatory activity was detected for seven, indicating that either the index or coding proxy variant could be causal (**Supplementary Figure 4I**). Similarly, haplotype-resolved WGS data revealed four instances in which other non-coding variants in high LD (r^2^ = 0.51–0.85) were likely responsible for the DE/ATU signals; these were labelled as “regulatory proxies” (**Supplementary Figure 4J**). We also identified ten variants whose regulatory effects are likely mediated in cell types or tissues distinct from those tested (“other tissue”, **Supplementary Figure 4K**), and seven where the regulated genes were not plausible candidates for the GWAS phenotype (“other gene”).

Overall, our curation yielded a high-confidence set of 22 variants for which we can propose strong mechanistic links between the regulated genes and blood traits (labelled “GeneBass, Open Targets or literature support”; **Figure 3E, Table 1** and **Table S6**). Among these, eleven involved DE events (**Supplementary Figure 4A–C**), seven involved ATU events (**Supplementary Figure 4D–F**), and 4 showed both (**Supplementary Figure 4G–H**). Notably, five variants had been previously reported as eQTLs for the same genes in the eQTLGen^4^ and/or BLUEPRINT^3^ consortia and one as a sQTL in the INTERVAL RNA-seq study^29^ (**Table 1)**.

**Table 1.**
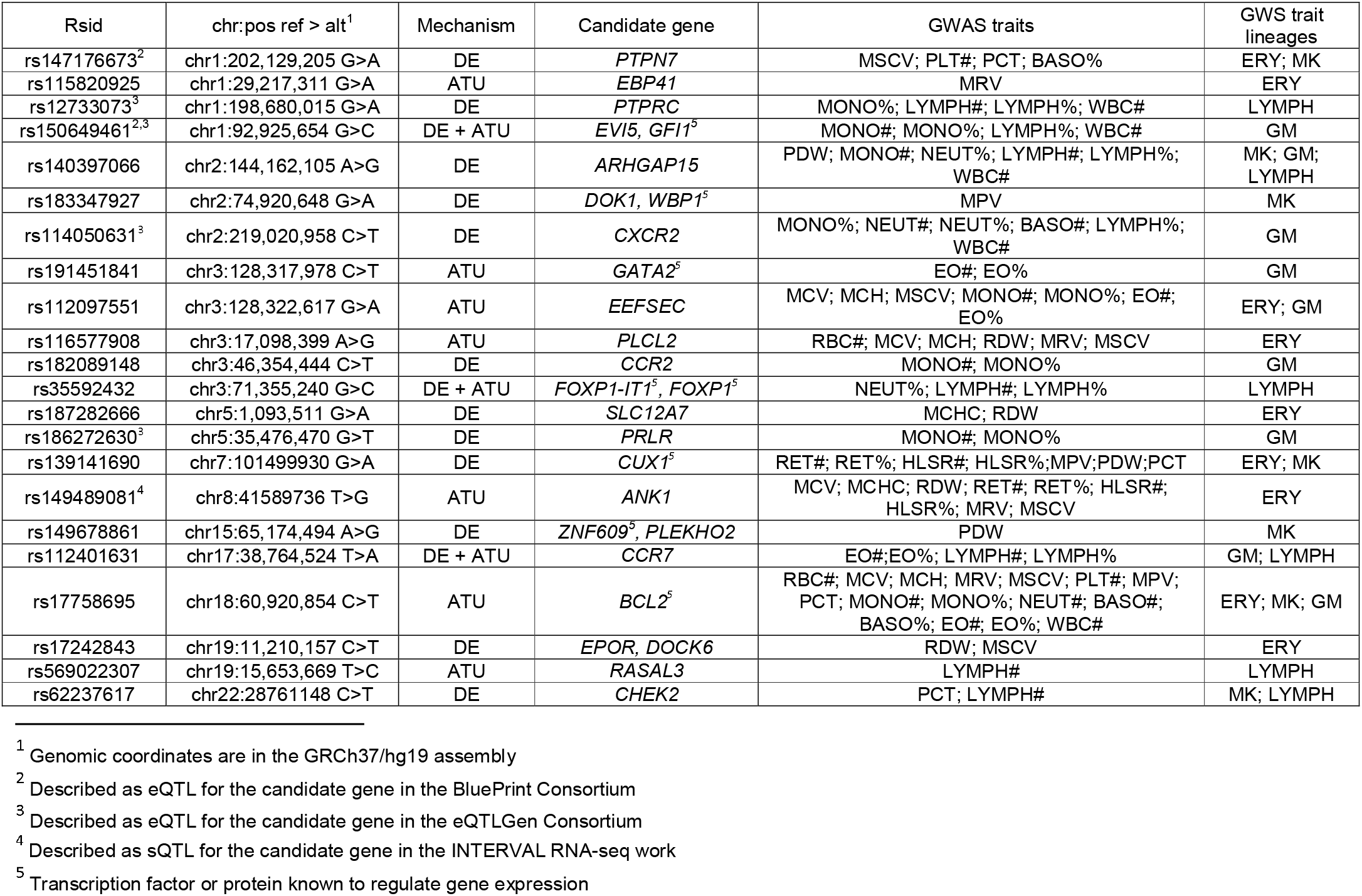
Variants with curated mechanistic hypothesis. *For the abbreviations of blood phenotypes see Table S3. DE = Differential Expression, ATU = Alternative Transcript Usage. MK =* megakaryocytic; ERY = erythroid; LYMPH = lymphocyte; GM = granulocyte monocyte. **Trait pleiotropy: No = Lineage restricted, Yes = Multi lineage**

### Regulation of megakaryocytic size and maturation by *cis* variation in *CUX1*

We selected four variants from the high-confidence set of 22 for experimental validation: rs139141690 (DE in the transcription factor *CUX1*), rs182089148 (DE in *CCR1* and *CCR2*), rs17758695 (ATU in *BCL2*), and rs35592432 (DE in *FOXP1-IT1* and ATU in the transcription factor *FOXP1*) (**Figure 4A, Supplementary Figure 4B, E**, and **H**). Three variants (rs139141690, rs182089148, and rs17758695) were MPRA positive, while rs35592432 marginally failed to reach the log_2_FC threshold.

**Figure 4.**
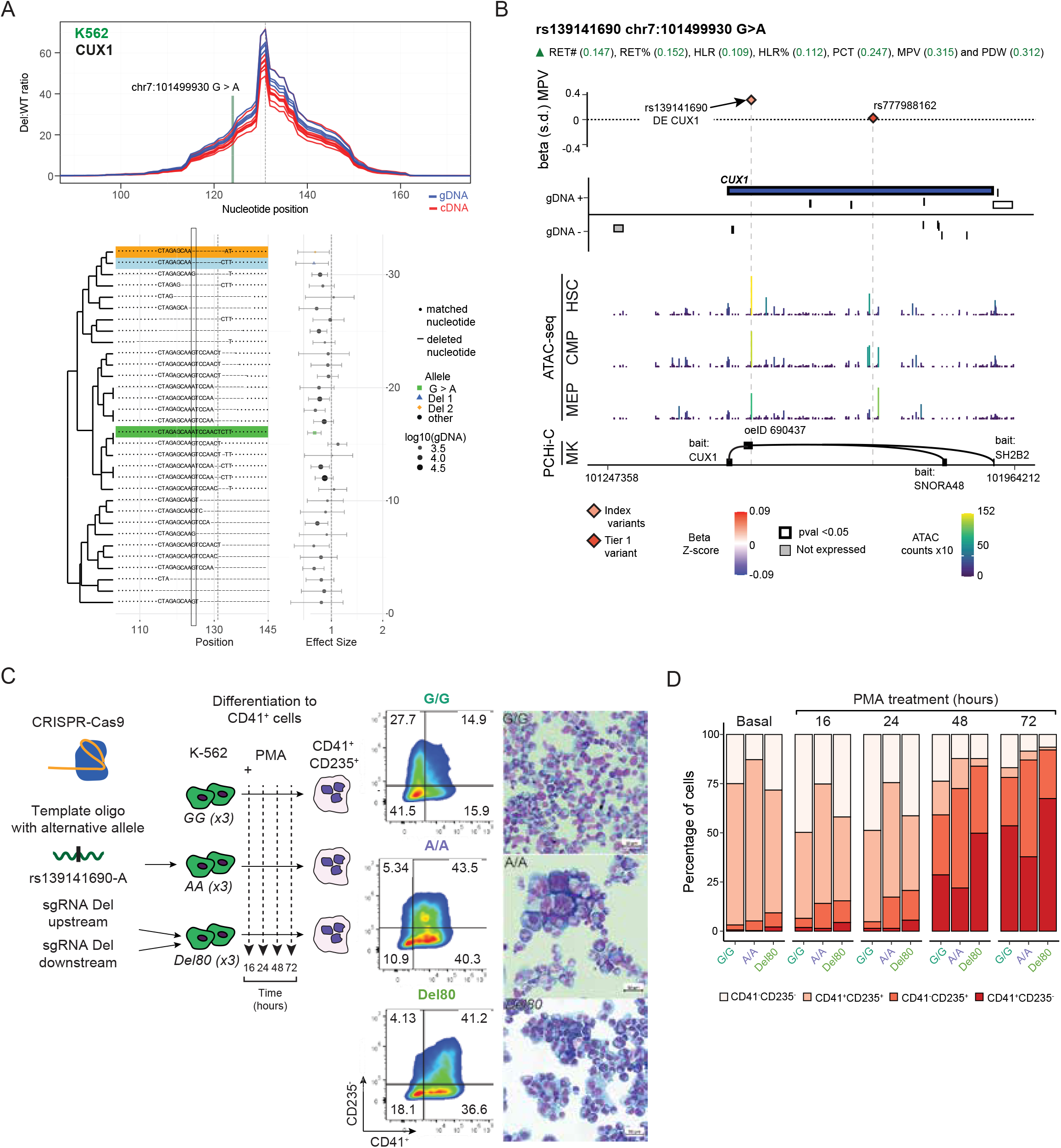
rs139141690-A downregulates the hematopoietic TF *CUX1*. **A)** GenIE results for the *knock-in* (highlighted in green) and UDPs (rest of profiles, two representative ones highlighted in blue and orange) of rs139141690 in K-562 cells. In blue the signal from genomic DNA and in red the signal from retrotranscribed pre-spliced mRNA. The highlighted UDPs and the *knock-in* allele significantly decreased the abundance of the mRNA in the edited cells. **B)** Locus plot for the variant rs139141690 that downregulates *CUX1* in whole blood. The variant sits in an intron of *CUX1* and has PCHi-C and accessible chromatin evidence in the megakaryocytic and erythroid lineages. **C)** Time course differentiation of the edited K-562 cells to CD41^+^ cells. **D)** Variation in the abundance of the different populations in the differentiation. The percentages represent the mean for the observation of the three clones per genotype. *See Table S1 for all abbreviations*.

CRISPR/Cas9 *knock-in* successfully generated clonal lines for rs139141690 and rs35592432. We then used the GenIE (Genome engineering-based Interrogation of Enhancers) screen^12^ to assess the effects of alternative alleles and unique deletion profiles (UDPs) on target gene expression. The rs139141690-A allele significantly reduced *CUX1* transcript abundance in K-562 cells (**Figure 4A**), while deletions spanning rs35592432 significantly increased *FOXP1* mRNA levels in the human induced pluripotent cell line Kolf2 (**Supplementary Figure 5A**), supporting the *in vivo* regulatory effects of both variants.

To validate the proposed mechanisms, we next generated cellular models. For rs35592432, we edited H1 human embryonic stem cells (H1 hESCs) and differentiated them into monocytes, as ATU of *FOXP1* was observed in this lineage (**Supplementary Figure 5B**). Bulk RNA-seq revealed increased levels of a *FOXP1* transcript without a CDS and a significant enrichment of *FOXP1* target genes among DE genes (**Supplementary Figure 5C–D**), supporting the proposed mechanism of action.

For rs139141690–*CUX1*, preliminary data and genomic annotations indicated an enhancer element influencing *CUX1* expression in the megakaryocytic lineage (**Figure 4B, Supplementary Figure 6A–B, Methods**). We investigated its role by differentiating K-562 cells into megakaryocyte-like CD41^+^ cells (*ITGA2B*, a megakaryocyte marker) using phorbol myristate acetate (PMA)^31^. Three isogenic K-562 lines were engineered: reference (G/G), homozygous alternative (A/A), and an 80 bp deletion spanning the SNP and all significantly active UDPs from the GenIE assay (80 bp del) (**Figure 4C**). Flow cytometry for CD235 (*GYPA*, erythroid marker) and CD41 revealed that A/A clones accumulated more intermediate CD235^+^CD41^+^ cells but fewer mature single-positive CD41^+^ cells, whereas 80 bp del clones reached higher proportions of CD41^+^ cells more rapidly than other genotypes (**Figure 4D, Methods**). These findings confirmed that rs139141690 affects megakaryocyte differentiation *in vitro*.

We next performed single-cell RNA and open chromatin profiling (scRNA-seq/scATAC-seq) during K-562 megakaryocyte differentiation, including the above clones plus a heterozygous (A/G) clone and a shorter deletion (16 bp del) spanning the PU.1 binding site affected by the variant. A total of 11,250 cells were genotyped, clustering into 13 groups (**Figure 5A–B**). We focused on cluster 1 (late-stage megakaryocyte maturation) and cluster 3 (cells co-expressing megakaryocytic and erythroid markers) (**Figure 5B–C**).

**Figure 5.**
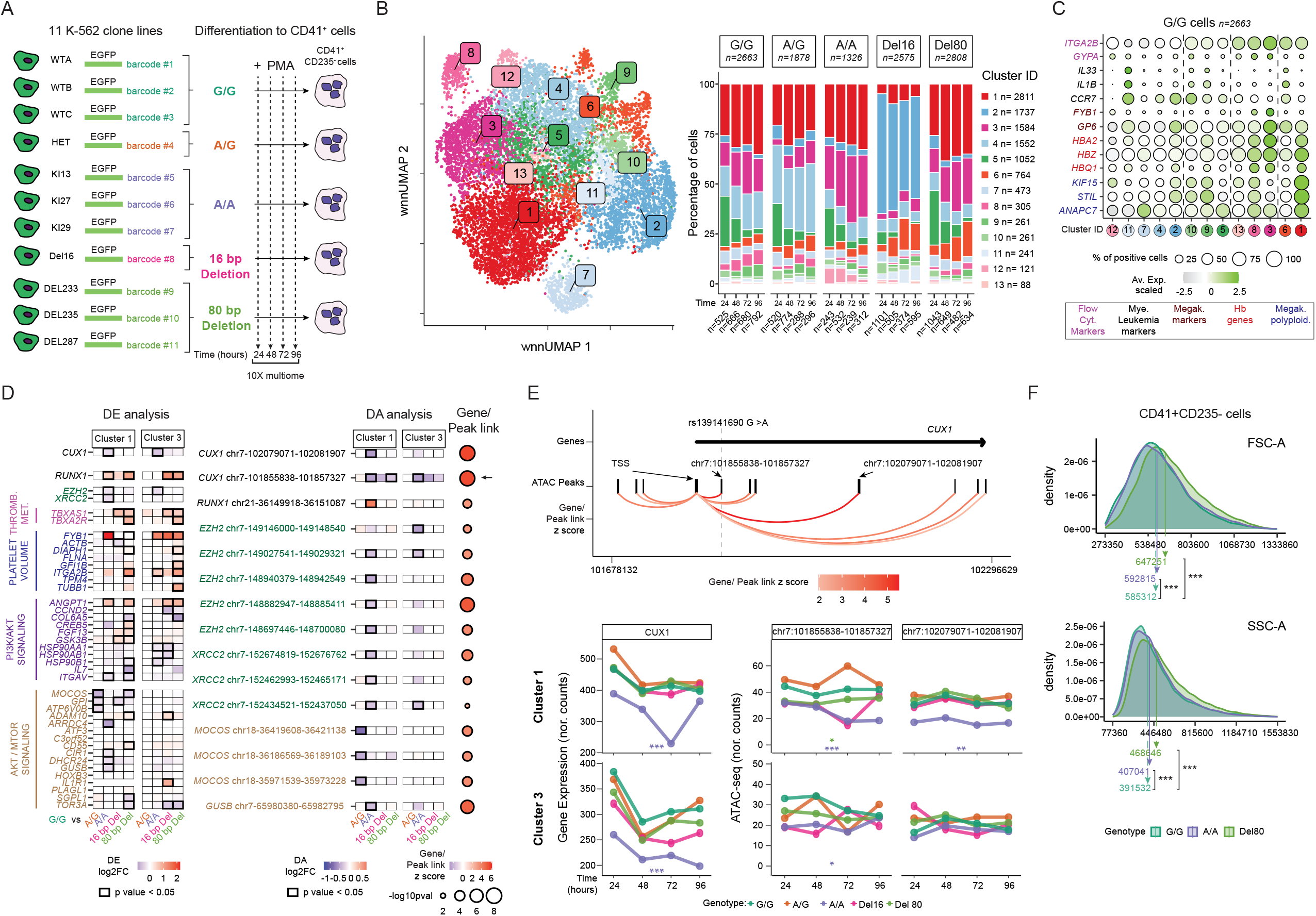
Single Cell multimodal analysis confirms the effects on gene expression and chromatin accessibility elicited by rs139141690-A that reveal a regulatory region with key effects on platelet size. **A)** Experimental design including the additional clone lines. **B)** U-MAP combining both sc-RNA-seq and sc-ATACseq modalities. To the right, changes in their relative abundance across time points and genotypes **C)** Expression of five sets of marker genes characterising key stages in the differentiation. The dashed lines indicate the separation of the clusters into 4 groups that we hypothesise correspond to the flow cytometry subgroups. **D)** DE and DA analysis in the clusters 3 and 1 for the comparisons of each genotype against wild type cells. The arrow indicates the peak overlapping rs139141690. **E)**. Locus plot detailing the DE and DA changes in clusters number 1 and 3 for the *CUX1* gene. **F)** FSC-A and SSC-A analysis across the genotypes in K-562 CD41^+^CD235^-^ cells at 72 hours. *Cyt. = cytometry, Mye. = myeloid, Megak. = megakaryocyte, Hb = Haemoglobin, polyploid = polyploidization, Thromb. Met. = Thromboxane metabolism, nor. counts = normalised counts*.

We analyzed the impact of each mutation relative to G/G cells in gene expression (DE) and chromatin accessibility (DA) (**Methods**). The heterozygous A/G genotype exhibited three DE genes and three DA peaks linked to the *AKT/mTOR* signaling pathway, consistent with a role of *CUX1* as a tumor suppressor^34^ gene (**Figure 5D)**. The A/A genotype showed broader dysregulation, including multiple DE genes within the PI3K/AKT/mTOR pathway, upregulation of *EZH2*, a polyploidization gatekeeper^35^, its target *XRCC2*, and increased expression of *FYB1*, implicated in abnormal platelet size^36^. DA analysis revealed significant accessibility changes near *EZH2* and *XRCC2*, consistent with the accumulation of CD235^+^CD41^+^ cells that fail to mature fully in the A/A genotype (**Figure 5D**). The two deletions produced complementary patterns: the long deletion (80 bp del) affected genes involved in megakaryocyte fate (*TBXAS1, TBXA2R*) and platelet volume (*ITGA2B, TUBB1*), whereas the short deletion (16 bp del) yielded fewer DE genes with concordant expression effects (**Figure 5D**).

To identify key transcription factors driving these changes, we focused on *RUNX1* and *CUX1*, both critical regulators of megakaryopoiesis^37^. *RUNX1* was significantly upregulated in the A/A and both deletion genotypes, but only A/A showed strong *CUX1* downregulation and reduced chromatin accessibility at the variant locus (**Figure 5D–E**). We assessed motif accessibility using chromVAR deviation scores^38^ for peaks containing *CUX1* and *RUNX1* motifs in clusters 1 and 3 (Methods). Deviation scores for *CUX1* motifs differed significantly from G/G only in the A/A genotype, whereas *RUNX1* motifs were significantly altered in A/A and both deletions (**Supplementary Figure 6C**). Gene regulatory network (GRN) analysis showed that the *CUX1* GRN was enriched among DE genes exclusively in A/A, while the *RUNX1* GRN was enriched in both A/A and the deletions (**Supplementary Figure 6D–E**), demonstrating the specificity of *CUX1* downregulation to the A/A genotype.

Finally, both 80 bp del and A/A cells produced significantly larger CD41^+^ cells after 72 hours compared to ‘G/G’ (p < 0.0001, Wilcoxon test; Figure 5F), linking the observed *in vitro* phenotypes with the increased mean platelet volume reported in the GWAS.

## Discussion

A previous comprehensive association analysis of blood indices by our group discovered 16,900 conditionally independent trait-variant associations^2^. Here, we leveraged these GWAS parameters to select 94 rare non-coding variants (RNVs) for functional characterization. RNVs represent an underexplored regulatory space, which we show to be independent of common alleles (in contrast to observations from other rare variant prioritization strategies^39^) and as having larger average effect sizes than reported pathogenic variants. A better understanding of the association properties and functional relevance of these variants sheds light into an unexplored but potentially clinically relevant set of human variation^6,8^.

We implemented a dual strategy integrating MPRA data on enhancer activity with differential expression and alternative transcript usage analyses from large RNA-seq datasets containing carriers for most index variants. A high proportion of variants (45.7%) showed evidence of enhancer activity and/or allelic skews from MPRA assays, aligning with prior evidence that these putatively causal variants with strong statistical support are indeed enriched for functional effects^10^. By jointly interpreting MPRA and RNA-seq results and conducting manual curation to exclude cases driven by proxy variants or unrelated genes, we identified 22 index variants with strong mechanistic support—16 of which lacked prior *in vivo* evidence of regulatory function. Notably, 12 of these 22 were MPRA^+^, while none of the variants identified as proxy of causal variants showed MPRA activity; this underscores the assay’s ability to capture biologically relevant effects.

This curation process also highlighted two limitations. First, MPRA-negative variants with RNA-seq evidence tended to overlap motifs for transcription factors expressed at low levels in each MPRA cell type, underscoring the need to expand future assays to additional cellular contexts. Second, despite the large scale of the bulk RNA-seq resources employed, statistical power for DE/ATU detection remained limited for variants with very low allele frequencies.

We provided *in vitro* experimental validation for two variants—rs35592432 and rs139141690— supporting the proposed regulatory mechanisms. For rs139141690, we demonstrated a causal link between *CUX1* expression and chromatin accessibility in the affected region. We propose that the observed increase in CD41^+^ cell size results from a combined effect of *CUX1* and *RUNX1* downregulation in the A/A genotype, with a stronger *RUNX1*-driven effect in the 80 bp del genotype. Notably, rs139141690 was recently included in an independent CRISPRi screen that confirmed *CUX1* downregulation in the targeted region but did not detect a variant-specific effect in K-562 cells^13^. This discrepancy may reflect methodological differences: CRISPRi assays infer editing indirectly via sgRNA detection and lack direct genotypic resolution, which may increase false negatives. In contrast, our approach—using fully genotyped clones followed by lineage-specific differentiation—provides higher sensitivity for detecting allele-specific regulatory effects.

Overall, this study provides proof-of-principle evidence for the value of systematic frameworks to prioritize and experimentally assess rare non-coding variants (RNVs), with applicability extending well beyond the context of blood traits. We anticipate that such efforts will be further enabled by improved fine-mapping of large-scale GWAS and by the expansion of molecular profiling datasets to encompass thousands of donors, ideally at single-cell resolution. Overcoming these challenges will be essential for generating mechanistic insights into the target specificity and regulatory modulation mediated by rare non-coding variants.

## Acknowledgements

<u>Acknowledgements</u>: Tristram Bellerby, Mathew Mayho, Jeremy Schwartzentruber, Sarah Cooper, Andrew Bassett from the Wellcome Sanger Institute. Cecilia Dominguez-Conde, Davide Bolognini and Edoardo Giacopuzzi from the HT Population and Medical Genomics centre. Alessio Palini, Nicolò Panini, Silvia Bombelli in the HT Flow Cytometry Applications Resource unit and Clelia Peano, Eugenia Ricciardelli, Javier Cibella, Fabio Simeoni, Niccolò Alfano, Paolo Ferrari and Luigi Lamparelli in the HT Genomic Facility.

This work has been funded by BHF Cambridge Centre for Research Excellence RE/18/1/34212, the Chan Zuckerberg foundation and MIUR/MEF through Fondazione Human Technopole. M.I. and E.P. were supported by core funding from the British Heart Foundation (RG/18/13/33946: RG/F/23/110103), NIHR Cambridge Biomedical Research Centre (NIHR203312) [^*^], BHF Chair Award (CH/12/2/29428), and by Health Data Research UK, which is funded by the UK Medical Research Council, Engineering and Physical Sciences Research Council, Economic and Social Research Council, Department of Health and Social Care (England), Chief Scientist Office of the Scottish Government Health and Social Care Directorates, Health and Social Care Research and Development Division (Welsh Government), Public Health Agency (Northern Ireland), British Heart Foundation and the Wellcome Trust. M.I. was also supported by the UK Economic and Social Research 878 Council (ES/T013192/1). ^*^The views expressed are those of the authors and not necessarily those of the NIHR or the Department of Health and Social Care.

RNA-seq in the INTERVAL study was funded as part of an alliance between the University of Cambridge and the AstraZeneca Centre for Genomics Research, and by the NIHR Cambridge Biomedical Research Centre (BRC-1215-20014). Participants in the INTERVAL randomised controlled trial were recruited with the active collaboration of NHS Blood and Transplant England (www.nhsbt.nhs.uk), which has supported field work and other elements of the trial. DNA extraction and genotyping were co-funded by the National Institute for Health and Care Research (NIHR), the NIHR BioResource (http://bioresource.nihr.ac.uk) and the NIHR Cambridge Biomedical Research Centre (BRC-1215-20014) [^*^]. The academic coordinating centre for INTERVAL was supported by core funding from the: NIHR Blood and Transplant Research Unit (BTRU) in Donor Health and Genomics (NIHR BTRU-2014-10024), NIHR BTRU in Donor Health and Behaviour (NIHR203337), UK Medical Research Council (MR/L003120/1), British Heart Foundation (SP/09/002; RG/13/13/30194; RG/18/13/33946), NIHR Cambridge BRC (BRC-1215-20014; NIHR203312) [^*^], and by Health Data Research UK, which is funded by the UK Medical Research Council, Engineering and Physical Sciences Research Council, Economic and Social Research Council, Department of Health and Social Care (England), Chief Scientist Office of the Scottish Government Health and Social Care Directorates, Health and Social Care Research and Development Division (Welsh Government), Public Health Agency (Northern Ireland), British Heart Foundation and Wellcome. A complete list of the investigators and contributors to the INTERVAL trial is provided in Di Angelantonio et al^40^. The academic coordinating centre would like to thank blood donor centre staff and blood donors for participating in the INTERVAL trial.

^*^The views expressed are those of the authors and not necessarily those of the NIHR or the Department of Health and Social Care.

## STAR methods

### Experimental model and study participants

#### MPRA data

The MPRA raw fastq files will be uploaded to the European Nucleotide Archive (ENA) upon publication.

#### ENCODE K-562 expression data

We used processed RNA-seq count matrices from basal K-562 cells.^41^

#### INTERVAL and BluePrint data

The INTERVAL study data used in this paper are available to bona fide researchers from. The data access policy for the data is available at http://www.donorhealth-btru.nihr.ac.uk/project/bioresource. The RNA-seq data in the INTERVAL cohort have been deposited at the European Genome-phenome Archive (EGA) under the accession number EGAD00001008015 and are available at^42^. The UK Biobank genetic data used in this study were approved under application 82779 and are available to qualified researchers via the UK Biobank data access process. For the BluePrint data we used data from^5^. All data are freely available but managed by the BLUEPRINT Data Access Committee.

#### 10X multiome data

The 10X multiome raw fastq files will be uploaded to the European Nucleotide Archive (ENA) upon publication.

#### Cell culture

K-562 (ATCC® CCL-243™, sex female), and HL60 (ATCC®CCL-240, sex female) cells were cultured as indicated by the distributor, 1x RPMI 1640 media with L-glutamine (Gibco Medium 52400025), supplemented with 10% Foetal Bovine Serum (FBS) (Gibco, A31604-02) and 1x penicillin/streptomycin (Gibco, 15070-063). THP-1 (ATCC® TIB-202, sex male) were cultured as indicated by the distributor, 1x RPMI 1640 media with L-glutamine, 2-mercaptoethanol (Sigma, M3148) was added to a final concentration of 0.05mM and supplemented with 10% FBS (Gibco, A31604-02) and 1x penicillin/streptomycin. CHRF-288-11 (sex male, a kind gift from Prof. Wilen H Ouwehand’s lab) were cultured 1x RPMI 1640 media with L-glutamine, supplemented with 20% Horse Serum (Gibco 16050-122) and 1x penicillin/streptomycin. All the cell types were maintained up to a confluence of 1x10^6^ cells/milliliter (ml) and then reseeded at 1x10^5^ cells/ml, except for THP-1 that were reseeded at 3x10^5^ and cultured in a T75 flask in an upright position. Phoenix Ampho (ATCC, CRL-3213, sex female) cells were cultured in DMEM 10% FBS prior to viral infections.

Induced Pluripotent Stem Cell (iPSC) line Kolf2_c1 line (Wellcome Sanger Institute’s Human Induced Pluripotent Stem Cell Initiative, sex male) was cultured in TeSR-E8 complete culture media (Stem Cell Technologies #05991) (37°C, 5% CO2) on 10 ng/ml Synthemax-II (Corning CLS3535) coated plates. Kolf2_c1 were thawed into TeSR-E8 + 10% CloneR (StemCell Technologies # 05888) and split at 70-80% confluence into TeSR-E8 + 10 micromolar (uM) Y27632 Rock Inhibitor (StemCell Technologies #72302).

H1 embryonic stem cells are cultured in Essential 8 medium (Gibco) supplemented with penicillin streptomycin in wells coated with BG iMatrix-511 (Tebu Bio).

At defrosting, for CRISPR/Cas9 editing and clones isolation or when cultured from single cells, H1 were cultured in Stem Flex media (Gibco) supplemented with penicillin streptomycin.

To differentiate K-562 cells to CD41^+^ cells, we seeded 100.000 cells/ml in IMDM + 10% FBS and cultured in presence of 5 nanomolar (nM) phorbol 12-myristate 13-acetate (PMA, Selleckchem) or DMSO for 16, 24, 48 and 72 hours.

## Method details

### Variant Annotation

The 178,890 variants in the 95% credible sets of the 29 blood indices from Vuckovic et al^2^ were prioritised attending to MAF in UK Biobank (1% threshold), Variant-Effect-Predictor (VEP)^43^ Most Severe Consequence (MSC), Posterior Probability (PP) and effect size (scaled to amount of standard deviation units per trait^2^). The prioritised subset of index variants were below or equal 1% MAF, MSC non-coding, with at least one blood index association above or equal 0.9 PP and with an effect size for that association between the absolute minimum and first quartile (beta < Q1) or the third quartile and the absolute maximum (beta > Q3), **Figure 1A** and **Table S2**. We condensed the VEP MSC option^44^ into coding and non-coding consequences. The coding group comprised the following labels: LOF (<u>L</u>oss <u>O</u>f <u>F</u>unction, [splice_acceptor_variant, splice_donor_variant, stop_gained and frameshift_variant]), MISS (missense_variant), UTR5 (5_prime_UTR_variant), UTR3 (3_prime_UTR_variant) and SYN (synonymous_variant). The non-coding group comprised the following labels: INTRON (intron_variant), INTERGENIC (intergenic_variant), UPSTREAM (upstream_gene_variant), DOWNSTREAM (downstream_gene_variant), REGULATORY (regulatory_region_variant), TFBS (TF_binding_site_variant), SPLICE (splice_region_variant), OTHER (start_lost, stop_lost, inframe_deletion, inframe_insertion, stop_retained_variant and mature_miRNA_variant), NMD (NMD_transcript_variant) and NCT (non_coding_transcript_variant). PCHi-C data was downloaded from Javierre et al^45^ (PCHiC_peak_matrix_cutoff5.tsv). ATAC-seq data^46^ was for blood cell types was downloaded from^47^ (29August2017_EJCsamples_allReads_500bp.bed and 29August2017_EJCsamples_allReads_500bp.counts.txt) and intersected with our variants.

An updated fine-mapping analysis in the Apr 2023 release to the GWAS catalog revealed 29 more index-variants meeting the inclusion criteria. These additional variants could not be included in the MPRA experiment, which was by then ongoing. We annotate them for completeness in **Table S2** (denoted in the Observations field with *“*^*\*\**^ *found after updating parameters for the GWAS catalog release”*) but remove them from further analyses for the sake of clarity.

### MPRA Library design and cloning

We designed a library of 19,050 200-mer oligonucleotides that were synthesised by Twist Bioscience. The library covered 113 SNPs, each one assayed in five partially overlapping tiles, every tile having an alternative and reference allele version. Each reference or alternative allele tile was tagged by 15 unique 11 bp barcodes. The library included 7 enhancer and allelic skew positive controls and 8 enhancer positive controls from^46^ and four sequences that showed no CRISPRa activity in Fulco et al^22^ as negative controls. The structure of the 200-mers included two 15 bp amplification arms at each end to amplify subpools/bins of the library based in GC content, an 11 bp barcode, the restriction enzyme sites for *BamH*I and *KpnI* and 148 bp of candidate regulatory sequence to be assayed. The amplification PCRs (Primers 1-6 **Table S7**) were done with Kapa HiFi HS Ready Mix (Kapa Biosystems), using 20 ng input template and 50 ul final volume with the exception of the High GC bin in which 20 microliters (ul) of KAPA2G GC Buffer (ROCHE) were added to a final volume of 100 ul. Next we performed a digestion with *ExoI* (NEB) to eliminate free primers and then purified the amplified fragments using Agencourt AMPure beads (Beckman Coulter).

We based the backbone vector for this assay on the hSTARR-seq_ORI vector^15^ (Addgene #99296) following the recommendations from Muerdter et al^15^. We added GFP to the vector by excising sgGFP from the pSTARR-seq_human vector (Addgene Plasmid #71509^48^) with an o.n. digestion with *AflII* and *Age*I (New England Biolabs, NEB) at 37°C and ligating it with hSTARR-seq_ORI vector digested in the same manner (T4 DNA ligase 16E o.n. 3 to 1 molar ratio). Following an idea proposed in the Supplementary Figure 1 I) of Muerdter et al^15^, we excised the polyA site from the hSTARR-seq_ORI GFP vector to be cloned back at a later stage of our library construction to separate the barcode and the candidate regulatory sequence. The excision was carried out by *Nae*I digestion of the vector (60’ 10 Units of enzyme, NEB) and posterior blunt end ligation to obtain the hSTARR-seq_ORI GFP polyA MINUS vector. The excised SV40 poly (A) signal fragment was amplified (Primers 9-10 **Table S7**) and cloned in the pGEM T easy system (Promega).

The amplified subpools of the library and the hSTARR-seq_ORI GFP polyA MINUS vector were ligated using Gibson cloning (NEB). Briefly, the vector was linearized by PCR (Primers 7-8 **Table S7**) and subsequently we carried out a digestion with *DpnI* and *BamH*I (NEB) to degrade the circular template. The ligation in Gibson mix was done with 100 ng of the linearized vector and a molar insert:vector ratio of 2:1 for each of the bins. The ligations were purified with Agencourt AMPure beads and eluted in 20 ul of Elution buffer diluted 1/10 in nuclease-free water and 10 ul of each were then used to electroporate electrocompetent E. coli bacteria (NEB) at 2000 volts 25 uF and 200 Ohm. We performed serial dilutions for each of the bins to ascertain the yield in colony-forming units (CFU) and aimed to keep a ratio of at least 100 CFU per oligo element. The individual colonies in each bin were lysed and the plasmids corresponding to each bin were purified using the Qiagen maxi prep kit (cat. 12162). This intermediate step in the library construction (PolyA Minus library) was sequenced to check the barcode - candidate regulatory sequence association and the design dropout rate. Briefly, we amplified 25 ng of each of the poly (A) minus library bins (Medium GC, High GC and Low GC content) with oligos 11 and 19 (**Table S7**) for 15 cycles with annealing and extension done at 72°C in a combined step for 1’. The libraries were then quantified with KAPA Illumina SYBR Universal Lib Q. Kit. (Roche), adjusted at 4nM and pooled together for a final volume of 40 ul. The samples were subsequently sequenced in a MiSeq (MiSeq Reagent Kit v2 300 cycle, Illumina) with the first 15 cycles of read 1 set to dark cycling and using custom primers 22 and 23 (**Table S7**). The PhiX amount was set to 10%.

Finally the polyA site was introduced back into the vector separating the barcode from the regulatory sequence using the *BamH*I and *KpnI* restriction sites *t*hat were introduced in the oligo design. We digested the PolyA Minus library with *Kpn*I (NEB, 37°C o.n.), followed by gel purification and digestion with *BamHI* in the presence of Shrimp Phosphatase (NEB) for 2h at 37°C. In parallel, we digested overnight 20 ugr of the pGEMT-polyA vector to release the SV40 polyA signal (230 bp) with *KpnI* and *BamHI* and gel purified it. We cleaned both fragments with Agencourt AMPure beads and quantified them using Qubit 1X dsDNA BR Assay Kit (ThermoFisher). For each bin we used 100 ng of the input vector and a 2:1 insert:vector ratio in the T4 ligase 16°C overnight reactions. We purified the ligations with Agencourt AMPure beads and performed electroporations in the same conditions as the ones used for the PolyA Minus library. After evaluating the efficiency of the electroporation we seeded 0.5 million CFU per 245 mm Square BioAssayDish with Agar+Ampicillin, keeping at least 100 CFU per oligo element. A step-by-step protocol of the procedure is available in protocols.io^49^.

### MPRA Nucleofection and parallel mRNA and gDNA isolation

Cells were seeded at 1.5 to 2x10^5^ cells/ml into 2-4 T175 flasks (Corning, CLS431085-50EA) in 60 ml of fresh medium (9 to 12 million cells each flask), and cultured for 48 hours changing half of their media 24 hours after seeding. The electroporation was performed using the Neon Transfection System (ThermoFisher). In K-562 and HL-60, we electroporated 5 million cells with 25 ugr of a pool of the three plasmid bins per reaction (12.5 ugr of the Medium GC bin and 6.25 ugr of each of the other two bins). In the case of CHRF-288-11 and THP-1 cells we electroporated 5 million cells with a four plasmid mix (10 ugr of the Medium GC bin, 5 ugr of each of the other two bins and 5 ugr of the pMAX-GFP vector). The nucleofection was performed following the manufacturer instructions using buffer R. The conditions for K-562, THP-1 and CHRF-288-11 cells were 1.45 volts, 10 milliseconds (ms) pulse width and 3 pulses, for HL60 1.350 volts, 35 ms pulse width and 1 pulse. The efficiency of the nucleofection was evaluated at 24h and 48h post nucleofection in the pmaxGFP plate with a Countess II FL Automated Cell Counter. We routinely observed 80-90% efficiency for K-562 cells. In the case of CHRF-288-11 and THP-1 due to the low transfection efficiency cells were co-transfected with pMAX-GFP and we used FACS to purify GFP positive cells enriching cells that have incorporated plasmids in the nucleofection step.

48 h post-electroporation cells were spun down (300G 5’) and resuspended in DNase I (NEB) at 37℃ for 15’ using 10 units (U) of enzyme per ug of plasmid transfected in a final volume of 2 ml of DPBS to eliminate possible carryover plasmid in the exterior of the cells. Cells were then pelleted (300 G for 5’), washed twice with DPBS and lysed in 600 ul of Buffer RLT Plus (Qiagen) with added β-mercaptoethanol and homogenised using QIAshredder columns (Qiagen 79654). DNA and total RNA were extracted using AllPrep DNA/RNA Kit (Qiagen) according to the manufacturer’s instructions. In the RNA preparation, a step of on-column DNase I treatment was performed for all samples (Rnase_Free Dnase Set, Qiagen). We isolated mRNA from total RNA using the Oligotex mRNA Kit (Qiagen) followed by a final treatment with Turbo Dnase (Invitrogen). DNA and mRNA quantifications were done using Qubit RNA Quantification, high sensitivity assay (ThermoFisher). A detailed protocol can be found in^50,51^.

### MPRA Library preparation and sequencing

We retrotranscribed 1-1,5 ug of mRNA per replica following the protocol for SuperScript IV (ThermoFisher) using a reporter specific RT primer (Primer 24, **Table S7**) at 2uM carrying the 10-mer UMIs. Then we split retrotranscription samples for PCR amplification so as the RT template would represent 10% of the final volume of the PCR (50 ul). We performed the first round of amplification in which we introduced the sample index primer, i5-i8 (Primers 15-18 **Table S7**) for the cDNA samples of four replicas. As a reverse primer, we used P7 (Primer 21 **Table S7**). The PCR was carried out using Kapa HiFi HS Ready Mix, and 65℃ annealing temperature for a total of 3 cycles. The amplification from each replica was then pooled, purified using Agencourt AMPure XP beads and then we assessed the minimum number of cycles for a second round of PCR by q-PCR with P5 and P7 with StepOnePlus™ Real-Time PCR System (ThermoFisher, Primers 20 and 21 **Table S7**). We determined between 11 and 13 cycles to be a good average range to keep the second round PCR from plateauing. Following Klein et al^23^ we split each of the replicas into 8 reactions and performed the second round PCR for the cycles determined with the P5 and P7 primers and Kapa HiFi HS Ready Mix at 64℃ annealing temperature. We then pooled the reactions from each replica, purified using Agencourt AMPure XP beads and eluted in 60 ul of Elution Buffer.

The gDNA fraction from the All Prep Qiagen kit was used as a source for the episomal plasmid nucleofected in every replica. For every replica, we used 12 ugr of gDNA that was split into 24 PCR reactions following Klein et al^23^. In this first reaction we introduced the sample index and the Unique Molecular Identifiers (UMIs) using the primers 11-14 (**Table S7**) as forward primers and primer 24 (**Table S7**) as reverse primer. The PCR was run for 3 cycles at 65℃ annealing temperature and the reactions corresponding to each replica were pooled, purified using Agencourt AMPure XPbeads and eluted in 320 ul of Elution Buffer. As in the cDNA library preparation we assessed by qPCR the number of cycles to keep the second round PCR from plateauing. We determined 10-11 cycles. We split each of the replicas into 29 reactions and performed the second round PCR with the P5 and P7 primers and Kapa HiFi HS Ready Mix at 72°C annealing temperature. Finally, we pooled the reactions for each replica, purified 200 ul of the mix using Agencourt AMPure XP system (1.2X vol/vol of beads) and eluted in 30 ul of Elution Buffer.

All the libraries were quantified using KAPA Illumina SYBR Universal Lib Q. Kit, adjusted to 4nM and pooled afterwards in 40 ul final volume. We used a HiSeq 2500 RR for sequencing each batch with the Hiseq Rapid PE Cluster Kit V2 (Illumina) and the HiSeq Rapid SBS Kit v2 200 cycles (Illumina FC-402-4021). The recipe included the first 15 cycles of read 1 set to dark cycling and used custom primers 22, 25 and 26 from **Table S7**. The amount of PhiX was set to 10%. A detailed protocol can be found in^51^.

### MPRA alignments and count matrix

The first six nucleotides of the r2 reads corresponding to the *BamHI* site were trimmed. UMIs were added to each read ID in r1 and r2 files using UMI tools^52^ and merged using flash^53^ prior to aligning against the reference set of barcodes using bwa^54^. Only primary alignments were taken forward. We discarded alignments not matching perfectly the corresponding barcodes with bamtools^55^ and deduplicated the UMIs per barcode using UMI tools. Finally we counted all the unique UMIs across all the tagging barcodes of a regulatory sequence.

### INTERVAL WGS analysis workflow

The INTERVAL whole genome sequencing data (WGS) were generated at the Wellcome Sanger Institute. The manuscript describing WGS in full is in preparation. Briefly, WGS was performed on 12,354 samples using the Illumina HiSeq X10 platform as paired-end 151 bp reads. Raw read processing was carried out via customised pipelines at WSI. Reads were aligned with BWA MEM to the GRCh38 human reference genome with decoys (also known as HS38DH). Variants were called for each sample using GATK HaplotypeCaller version 4.0.0. Then all samples were merged, and the combined samples genotyped using GATK4.0.10.1. GATK Variant Quality Score Recalibration (VQSR) was used to identify probable false positive calls by assigning quality score log-odds (VQSLOD) separately for SNPs and INDELs using GATK VariantRecalibrator (v4.0.10.1). Sample quality control removed 491 samples in total, including 77 samples with coverage below 12x, 134 samples with > 3% non-reference discordance (NRD), 118 samples with > 3% FreeMix (VerifyBamID2) score, 221 samples failing identity checks, 30 samples swapped, 40 samples failing sex checks, 39 duplicates and 9 samples with possible contamination. Genotypes with allele read balance > 0.1 for homozygous reference variants, < 0.9 for homozygous alternative variants or not between 0.2-0.8 for heterozygous variants were removed. Genotypes were also removed if the proportion of informative reads was < 0.9 or read depth > 100. We performed additional variant quality control and filtered out variants that failed to meet the following requirements: call rate per site > 95%, mean genotype quality (GQ) value > 20, Hardy-Weinberg equilibrium (HWE) p-value > 1 x 10^-6^ only for autosomes. All monomorphic variants with alternative allele count (AAC) = 0 were further removed, although we kept all monomorphic variants with reference allele count (RAC) = 0. For chrX and chrY we applied an additional step to correct allele counts and frequencies due to female and male samples accounting for diploidy/haploidy in the PAR and non-PAR regions. Finally, the WGS data set contains 116,382,870 variants (100,694,832 SNVs and 15,688,038 indels) including 6,637,420 (5.7%) multi-allelic sites across 11,863 participants.

### INTERVAL RNA-seq analysis workflow

The INTERVAL RNA-seq data were generated and processed as previously described^29^. We mapped the RNA-seq data to the GRCh38 reference and quantified read counts as described previously^56^ with the difference of using GENCODE v31 across 4,731 samples passing quality control. Globin genes, rRNA genes, and pseudogenes were removed. 19,841 genes were selected with > 0.5 CPM in at least 1% of the samples. Gene expression counts were converted to FPKM, trimmed mean of M-values (TMM) normalised and log_2_ transformed. We used the probabilistic estimation of expression residuals (PEER) method^57^, implemented in the R package peer v.1.0^58^, to correct for latent batch effects and other unknown confounders. 50 PEER factors were calculated with age, sex, BMI, and 19 blood cell traits included as covariates.

For transcript quantification we used Salmon v1.1.0^59^. The Salmon index was built against GRCh38 cDNA. R packages tximport v1.14.2, AnnotationHub v2.18.0, BiocFileCache v1.10.2, BiocGenerics v0.32.0 were applied to obtain various count matrices from these quantifications at the transcript or gene level. We subsetted to 4,731 samples passing RNA sequencing QC and corrected sample swaps. We focused on the transcripts of the 19,841 genes that passed gene QC. From these transcripts, we selected transcripts with TPM ≥ 0.1 in at least 20% samples. Subsequently, TPM values were TMM normalised and log2 transformed.

### GTEx and INTERVAL sQTLs

For the 20 genes with ATU in our survey of INTERVAL whole blood RNA-seq (*ATL1, NPRL3, VMP1, ELP5, KDSR, RASAL3, SLC11A1, MFSD2B, GATA2, EEFSEC, TAF8, IKZF1, PILRB, ANK1, GFI1B, EVI5, TYMP, ARSA, ODF3B* and *PILRB*) we queried the GTEx^60^ web portal (release v8) and found sQTLs in whole blood for *PILRB, SLC11A1, ARSA, ANK1, MFSD2B, TAF8, GFI1B* and *RASAL3*. The same query in the INTERVAL web^61^ yielded 16 genes: *MFSD2B, PILRB, TYMP, ANK1, ARSA, EVI5, NPRL3, VMP1, GFI1B, SLC11A1, IKZF1, EEFSEC, ODF3B, TAF8 GATA2* and *RASAL3*. We obtained all the proxy variants at R^2^=0.7 in the European subpopulations (EUR) in a window size of 0.5 Mb for all the sQTLs in these genes using LDLinkR^62^. None of the index variants leading to ATU were proxies of the GTEx sQTLs. Three of the index variants with ATU genes (rs149489081-*ANK1*, rs543594419-*TYMP, ODF3B* and *ARSA* and rs187715179-*GFI1B*) were sQTLs in the same genes in INTERVAL.

### Predicting TF motifs intersecting rs139141690 A > G

For the 94 variants screened in MPRA we selected a sequence stretch of 38 bp centred on the SNP and extracted the reference allele sequence. In the case of the 80-bp deletion we used the complete 80 bp nucleotide stretch. We then substituted the reference allele with the alternative allele in the position of the SNP to obtain the alternative allele version. We predicted TF motifs in both the reference and alternative allele versions of the 38 bp nucleotide stretch with gimme motifs^63^ with the reference databases HOMER^64^ and JASPAR^65^, the -c option set to 0.85 and the maximum number of TF motif per nucleotide stretch set to 20. To assess TF occupancy, we intersected the motifs with CHIP-seq data from whole blood present in the CHIP atlas database^66^. In the case of variant rs139141690 the occupancy of the PU.1 motif in the reference genome is supported by CHIP-seq data of PU.1 in 121 experiments in whole blood (ids: ERX626856, ERX626869, SRX093183, SRX093189, SRX100429, SRX100443, SRX100576, SRX10144602, SRX10144603, SRX1023790, SRX1023791, SRX1023792, SRX1023793, SRX103224, SRX1048461, SRX1089832, SRX1089833, SRX1127545, SRX12684447, SRX12684451, SRX1431740, SRX14869351, SRX14869352, SRX14869358, SRX14869359, SRX18154277, SRX18154278, SRX18154279, SRX18154280, SRX190299, SRX19553539, SRX20230007, SRX20230008, SRX20230009, SRX2268282, SRX2268283, SRX2268284, SRX2268285, SRX2268286, SRX2268287, SRX24542848, SRX24542849, SRX2770854, SRX2770855, SRX2770856, SRX2770857, SRX3824041, SRX3824042, SRX4001818, SRX4001819, SRX4001820, SRX4001821, SRX4001958, SRX4001959, SRX4001960, SRX4001961, SRX4484984, SRX475793, SRX475794, SRX5141098, SRX5141099, SRX5574342, SRX5574343, SRX5574345, SRX5574346, SRX5574348, SRX5574350, SRX5574352, SRX5574354, SRX5574355, SRX5574356, SRX5574357, SRX5574359, SRX5574361, SRX5574362, SRX5574363, SRX5574364, SRX5574365, SRX5574367, SRX5574369, SRX5574370, SRX5574373, SRX5574375, SRX5574376, SRX5574379, SRX5574381, SRX5574385, SRX5574387, SRX5574392, SRX5574446, SRX5574447, SRX5574448, SRX5574449, SRX5574450, SRX5574451, SRX5574452, SRX5574453, SRX5574457, SRX5574458, SRX5574459, SRX5574460, SRX5574461, SRX5574462, SRX5574463, SRX5574491, SRX5574492, SRX5574493, SRX5574494, SRX5574498, SRX5574499, SRX5574500, SRX5574505, SRX5574506, SRX5574507, SRX627428, SRX627430, SRX698188, SRX698189, SRX794057, SRX9029196, SRX9029197, SRX9029208, SRX9029209). The FOXM1 motif is supported by whole blood CHIP-seq data in 1 experiment (id: SRX190187).

### GenIE CRISPR/Cas9 targeting and amplicon design

We followed the protocol described in Cooper et al^12^. The Wellcome Sanger Institute Genome Editing browser (WGE)^67^, was used to choose CRISPR gRNAs with NGG PAM site within 20 bp of the SNP locus and with less than 1-3 mismatch off-target hits predicted. To introduce the SNP of interest a 100 bp repair template oligonucleotide was designed. As a positive control for cutting we used the sgRNA against ENSG00000178927/ CYBC1/ EROS (numbers 39-43 from **Table S7**).

Primers were designed to amplify <250 bp across the SNP of interest, 40-60% GC, Tm 56-65°C (NEB Tm Calculator) and with adaptor sequence tails for MiSeq Sequencing (See **Table S7**). Reverse transcriptase primers were designed downstream of the amplicon in the mRNA sequence (See **Table S7**).

### GenIE Nucleofection

Guide RNAs (IDT) were annealed to tracrRNA (IDT) in duplex buffer (IDT 1072570) at 95°C for 2 min and cooled slowly to RT. Nucleofection on Kolf2_c1 was carried out as previously described^68^ and recovered onto 4 ng/ul Synthemax-II (Corning CLS3535) coated 6 well plates. K-562 and HL-60 were nucleofected following Lonza’s protocols. THP1 were nucleofected following Lonza’s primary monocyte protocol. Cells were cultured for 1-2 weeks until confluent and snap frozen as 2-3 x10^6^ cell pellets.

### GenIE Library preparation and sequencing

Genomic DNA was extracted using DNA MagAttract HMW extraction kit (Qiagen), following standard instructions and eluted in 100 ul H20. Total RNA was extracted using Direct-zol RNA Miniprep Plus kit (Zymo), TURBO DNAse treated and run on a 2100 RNA Nano Chip in a Bioanalyser (Agilent). Gene specific RT primers were annealed using 2 ug RNA, 2 uM RT primer and 10 mM dNTPs, heated to 65°C for 5 min and placed on ice. cDNA synthesis was setup using half the annealed RNA, Superscript IV and RNasin (Promega) on ice and heated 50°C 10min, 55°C 10min, 60°C 10 min, 80°C 10min, 4°C hold.

Genomic DNA and whole RNA were amplified with PowerUP SYBR green master mix (Applied Biosystems A25742) or Q5 Hot Start polymerase (NEB), 10 uM adaptor sequence primers with 4 PCR replicates for gDNA and 8 PCR replicates for cDNA. PCR conditions for PowerUP SYBR reactions were as previously described^12^. PCR conditions for Q5 Amplicons: 98°C 30s, (98°C 10s, 57°C 20s, 72°C 20s)x30, 72°C 2min. Amplicons were barcoded using WTSI PCR barcoding primers, pooled, gel extracted using Minelute kit (Qiagen 28604) and quantified by qPCR (KAPA library quantification). The prepared library was loaded onto a MiSeq System (Illumina) at 4 nM with 20% PhiX using MiSeq Reagent Kit v2 300 cycles.

### Generation of clone lines: CRISPR/Cas9 sgRNA designs and protocol

To introduce the rs139141690 (G>A) mutation into K-562 cells, a synthetic crRNA was selected using CRISPOR^69^ (**Table S7**, number 46). An 81 bp ssODN carrying the mutation G>A was chemically made by IDT with 4 phosphorothioate bonds (**Table S7**, number 47). To delete the 80 bp region encompassing the rs139141690 two crRNAs were selected one upstream and one downstream of the desired SNP (**Table S7**, 48-49).

The crRNA and trans-activating crRNA (tracrRNA) were synthesised by IDT. An electroporation enhancer was also bought by IDT and resuspended at 100 µM in water. SgRNAs are made by combining 160 µM of crRNA and tracrRNA (1:1 v/v) to get a final concentration of 80 µM and incubated at 37C for 30 min.

To assemble the Cas9/sgRNA RNPs, the sgRNAs and electroporation enhancer (0.8:1 volume ratio) were first mixed and then 40 µM S.p. Cas9 Nuclease (IDT) at 1:1 v/v was added. This mixture was incubated at 37C for 15-30 min prior use.

Electroporation was performed using SF Cell Line 4D-Nucleofector™ X Kit (Lonza) according to manufacturer’s instructions. The kit / program for K-562 used is SF kit / FF-120.

For the *knock-in* experiment, 50 picomol (pmol) of the *RNP* was electroporated into K-562 cells together with 4µM ssODN as HDR template. 15 min after electroporation cells are incubated with a combination of 0.5µM Trichostatin A (TSA, Selleckem) and 1µM M3814 (Selleckem) to enhance the HDR as previously reported in Shy et al^70^. 24 hours post treatment drugs are removed, and fresh medium is added^70^. For the deletion of 80 bp, 50 pmol of each of the two RNPs was electroporated.

After 24h cells were single cell sorted in 96 wells plates using MoFlo Astrios (Beckman Coulter). Propidium Iodide (Sigma Aldrich) was added prior to analysis as a cell viability dye.

To introduce the rs35592432 (G>C) mutation into H1 cells, a synthetic crRNA was selected using CRISPOR. An 81bp ssODN carrying the mutation G>C was chemically made by IDT with 4 phosphorothioate bonds. The crRNA and trans-activating crRNA (tracrRNA) were synthesised by IDT. An electroporation enhancer was also bought by IDT and resuspended at 100 µM in water. SgRNAs were made by combining 160 µM of crRNA and tracrRNA (1:1 v/v) to get a final concentration of 80 µM and incubated at 37C for 30 min.

To assemble the Cas9/sgRNA RNPs, the sgRNAs and electroporation enhancer (0.8:1 volume ratio) were first mixed and then 40µM S.p. Cas9 Nuclease (IDT) at 1:1 v/v was added. This mixture was incubated at 37C for 15-30 min prior use. Electroporation was performed using P3 Primary cells 4D-Nucleofector™ X Kit (Lonza) according to manufacturer’s instructions with the electroporation program CB-150. 50 pmol of the RNP was electroporated into H1 cells together with 4µM ssODN as HDR template. 30 min before electroporation cells were incubated with a combination of ART558 3µM and AZD468 1µM (Selleckchem) to enhance the HDR. 24 hours post treatment drugs were removed and fresh medium was added.

72h-post transfection H1 colonies were mixed and expanded. 48 hours later single cells were detached using Accutase and plated using limiting dilution method in 96 well plates in presence of CEPT (200µM Chroman-1, 12mM Emricasan, 2.8mM trans-ISRIB, 1x Polyamine Supplement) to isolate single clones.

### Generation of clone lines: isolation of clones and genotyping by amplicon sequencing

K-562 clones were let to grow from single cells for 10-14 days, then genomic DNA was extracted with the QIAamp 96 DNA QIAcube HT Kit and quantified using the QuantiFluor® dsDNA System (Promega). 2-5 ng of DNA from every colony were used as template for targeted amplicon sequencing (**Table S7**, 44-45). Samples were then indexed with Nextera XT DNA Library Preparation Kit (Illumina), pooled and sequenced on NovaSeq 6000 (Illumina, 500 cycles, PE). Data were analysed by using CRISPResso2 v2.2.12^71^ and clones with the desired genotypes were expanded to generate the modified cell lines.

In the same nucleofection we isolated: i) three clone lines homozygous reference for the SNP rs139141690 chr7:101499930 (‘G/G’), ii) one heterozygous clone line chr7:101499930 (‘A/G’) and iii) three homozygous alternative clone lines chr7:101499930 (‘A/A’). In addition to this we isolated: i) one homozygous clone line carrying the deletion of 16 bp (chr7:101499917 CTCACTAGAGCAAGTCC > C) and ii) three homozygous clone lines carrying the 80 bp deletion (chr7:101499894 GTTAGTGACTTCAAAAGCTGTCCTCACTAGAGCAAGTCCAACTCTTCCTCTA GTTCTGATGACTTCACGGCAGCCAACTG > G). All the coordinates are in GRCh37.

H1 clones were grown in Stem Flex medium (Gibco) from single cells for 7-12 days, then genomic DNA was extracted with the QIAamp 96 DNA QIAcube HT Kit and quantified using the QuantiFluor dsDNA System (Promega). 5-10 ng of DNA from every colony was used as template for Sanger sequencing, performed by Eurofins Genomics. In the same nucleofection we isolated three clone lines homozygous wild type reference for the SNP rs35592432 (G/G clones), and three homozygous alternative clone lines rs35592432 (C/C clones).

### Generation of clone lines: lentiviral barcoding

Plasmids carrying single unique barcodes were isolated from the Larry Barcode Library V1^72^ (Addgene, #140025) characterised by whole plasmid sequencing (Eurofins) to identify the different barcodes and used to transfect Phoenix Ampho (ATCC, CRL-3213) cells, together with the pMD2.G^73^ (Addgene, #12259) and psPAX2^73^ (Addgene, #12260) lentiviral packaging vectors. 48h after the transfection, the viral supernatant was collected, filtered, and used to infect the K-562 clones with the different genotypes, in presence of 6ug/ml polybrene. GFP^+^ cells were sorted after 4 days and expanded to obtain 11 different cell lines (K-562-Larry), each one carrying a specific barcode, detectable both at gDNA and mRNA level.

### Flow cytometry tracking of CD41 expressing K-562 cells

K-562 cells were seeded at 100.000 cells/ml in IMDM + 10% FBS and cultured in presence of 5nM phorbol 12-myristate 13-acetate (PMA, Selleckchem) or DMSO for 16, 24, 48 and 72 hours. For each time point cells were analysed by flow cytometry and collected for RNA extraction.

For flow cytometry tracking, cells were washed in PBS and incubated with anti-CD235a BUV395 (Clone GA-R2, BD Biosciences) and anti-CD41 AF700 (clone HIP8, Biolegend) antibodies for 20 min at 4°C in PBS + 1% FCS + 2mM EDTA. Cells were then washed and acquired with the CytoFLEX (Beckman Coulter) flow cytometer. DAPI (Sigma Aldrich) was added prior to analysis as a cell viability dye. Data was analysed using the FlowJo software.

### H1 cells differentiation into monocytes and FACS

Three H1 rs35592432 G>C edited clone lines (3 x G/G reference clones and 3x C/C homozygous alternative clones) were differentiated into monocytes using the StemDiff Monocyte kit (StemCell Technologies) following the instructions of the manufacturer. After 18 days of differentiation, we sorted separately the six clone lines selecting for DAPI negative and CD11b+ (anti CD11b-APC, BioLegend) cells. 30.000 cells per sample were then processed for total RNA extraction using the PureLink RNA Micro Kit.

### Bulk RNA sequencing

SMART-Seq v4 PLUS Kit was used to generate cDNA libraries from differentiated monocytes total RNA. Libraries were then purified, quantified and their quality was assessed by TapeStation system. 3’ RNA sequencing was performed on Novaseq 6000 sequencer.

### 10X Multiome nuclei isolation

For each time point, cells were analysed by flow cytometry and 50.000 cells per genotype were pooled for nuclei isolation. Single nuclei were isolated using the Nuclei Prep Buffer (Zymo Research), counted and processed following the Chromium Single Cell Multiome ATAC + Gene Expression workflow^74^.

### 10X Multiome aligning and QC

10x genomics multiome data were processed using Cell Ranger ARC (2.0.2) using default parameters and the provided reference genome GRCh38-2020-A-2.0.0. Initial filtering steps were applied to the raw gene expression and peak (ATAC) matrices of each sample using functions from the Seurat and Signac (v5) packages^75,76^. Cells with <500 gene features were first removed and the package scDBfinder^77^ was used to mark doublets in both sc-RNA-seq and ATAC-seq data. Further, cells with < 1000 peak features, >10% mitochondrial reads, and multiplets marked by cellranger were removed. At this stage, doublets were additionally identified in the ATAC data using Amulet^78^. Prior to merging samples, ATAC matrices were rebuilt to reflect unique fragment counts in 5kb genomic windows instead of peaks, using a custom pipeline^79^, and sc-RNA-seq matrices were adjusted to remove ambient contamination using CellBender^80^. The merged Seurat object was then filtered to retain cells that could be unequivocally assigned to a lentiviral Larry barcode. To deconvolute samples based on the lentiviral barcodes, reads that did not map to the reference genome were extracted from the original cellranger genome + transcriptome alignments (gex_possorted_bam.bam) using samtools view -f 4, converted to fastq, and then re-mapped to a new reference containing the 11 possible GFP+barcodes transgenes. Multimappers were then removed (samtools view -q 5) to keep only the reads uniquely mapping to each GFP barcode. For each read, molecular barcode (UMI), cell barcode and Larry barcode were recorded. Only cell/Larry barcodes combinations supported by a minimum of 3 UMI were retained. Cell barcodes were assigned to a genotype only if one cell/Larry barcodes combination was found. The RNA and ATAC reductions from the merged and filtered Seurat object were then integrated using the Weighted Nearest Neighbour (WNN) analysis following the steps and recommended parameters in guidelines^81^. For clustering analysis on the integrated WNN graph, the Leiden algorithm with resolution 2 was used. At this stage, a low-quality cluster with lower read counts and higher mitochondrial content was removed, as well as doublets, which were previously marked as such by both Amulet and scDBfinder. After these final filters, we called ATAC peaks on the remaining cells with MACS2 (2.2.9.1) using the Signac callPeaks function and generated a new feature ATAC matrix with these peak coordinates, which replaced the window matrix. WNN analysis was then repeated as above, but with 0.5 resolution for the final clusters. The final object was composed of 13 clusters, 11.250 cells, ∼29.000 genes and ∼357.000 peaks.

## Quantification and statistical analysis

### Comparisons between sets of variants

In the case of the comparison with pathogenic variants, the list was obtained from Table 1 Vuckovic et al^2^ and restricted to GWAS traits associated with the pathogenic variants. Index variants had higher effect sizes to reported heterozygous blood ClinVar/HGMD pathogenic variants^2^ (median absolute effect size values 0.169 and 0.13 respectively, p-value = 0.007, *Wilcoxon test*)

For the comparisons shown in **Figure 1C** we used pairwise Wilcoxon tests to assess statistically significant differences between categories, applying multiple testing correction (Benjamini– Hochberg) across all the comparisons. Median values of MAF: Tier 1 0.285, Tier 2 0.013, Tier 3 0.018, Tier 4 0.016 and index variants 0.006.

Median values of PPFM: Tier 1 0.01, Tier 2 0.422, Tier 3 0.012, Tier 4 0.986 and index variants 0.999. Median values of absolute effect size: Tier 1 0.019, Tier 2 0.092, Tier 3 0.062, Tier 4 0.075 and index variants 0.152. Median values of CADD raw: Tier 1 -0.009, Tier 2 1.189, Tier 3 -0.049, Tier 4 0.034 and index variants 0.197. Median values of Gnocchi: Tier 1 0.653, Tier 2 1.834, Tier 3 0.868, Tier 4 1.203 and index variants 0.964. Median values of NCBoost: Tier 1 0.029, Tier 2 0.076, Tier 3 0.031, Tier 4 0.045 and index variants 0.079.

### Conditional analysis of common variants on the index variants

To condition for known common variants associated with the studied traits, we run GWAS of 29 blood cell counts in ∼409 UK Biobank individuals of white British ancestry, following the procedure described in^1^. Accordingly, we employed the same phenotype exclusions, adjustments and normalisation approach, and ran GWAS using REGENIE^82^ and TOPMed imputed genotypes^83^, including recruitment center and the first ten PCs of the kinship matrix as covariates. We then obtained independently associated variants for each trait by using the GCTA (v1.94.1) cojo^84^ joint model function (with parameters: collinearity = 0.9, p-value < 1e-4) and LD structure estimates from the genotypes of 30,000 unrelated individuals of white British ancestry from UK Biobank samples. Finally, for each of the 123 rare variants, we gathered all independent GWAS common variants obtained by cojo, having MAF > 1% and joint association p-value < 1x10^-7^, and falling within a +/- 500kb window from a rare variant. We then performed conditional analysis by fitting a linear regression model of the blood trait and the genotype of the associated rare variant, including the genotypes of the independent GWAS hits as covariates. The model was fitted in R (using the stats::lm function) and genotypes of both common and rare variants were obtained from the WGS of UK Biobank 200k release (GraphTyper population level WGS variants, PLINK format). Out of 123 rare non-coding variants, genotypes were available for 80 of them; of these 80, 76 (95%), were significantly (p-value < 0.05) associated with at least one blood trait even after jointly conditioning for common variants.

### MPRA analysis

We used MPRAmodel^24^ to estimate enhancer activity (measured as log2 Fold Change, log2FC) and allelic specific expression (measured as log2 Allelic Skew, log2AS) per each tile, index variant and cell type. First we generated the necessary input files for MPRAmodel per cell type: a count matrix with the oligos and the barcodes in the rows and the columns (countsData), a file detailing the replicate breakdown between cDNA and gDNA libraries (condData) and an attributes files detailing each of the oligos attributes (attributesData). In the majority of the cases the tiles carried only one SNP so the ‘Allele’ field of the attributes table was set to ref or alt. For the 8 cases of diplotypes (two SNPs present in the same tile) we carried out all the possible comparisons and set the ‘Allele’ field of the attributes table to ref ref vs alt ref, ref ref vs ref alt and ref ref vs alt alt according to the tiles analysed. Next we adapted the MPRAmodel Rscript^85^ from the MPRASuite to run the dataOut function on our inputs. The results were collected per cell type and tile and are shown in **Table S3**. We then performed a meta-analysis for each variant across all the tiles assayed to come up with a single value of activity per variant and cell type following^10,86^. The results of the meta-analysis are shown in **Table S4**. To establish the log2FC threshold that defines enhancer activity we employed a set of variants previously described to have MPRA activity in K-562 cells^9^ as well as four regions deprived of CRISPRa activity in the same cell line^22^(**Supplementary Figure 1A**). We considered active variants those with a log2FC higher than 0.25 at 1% global false-discovery rate (gFDR) (68 variants) and we additionally required that the log2AS was significant at 10% gFDR to label variants as MPRA positive (43 variants). Positive and negative Log2FC values indicate enhancer and repressor activity, respectively. Positive log2AS values correspond to a skew of the enhancer activity towards the alternative allele and negative values towards the reference allele.

For all the MPRA positive variants shared between two cell lines or more, we calculated the Pearson correlation coefficient for the values of Log2FC and Log2AS. The Pearson correlation coefficient values and the p-values were as follows:

K-562 vs CHRF: Log2FC correlation coefficient 0.736, p-value 0.0063.

K-562 vs CHRF: Log2AS correlation coefficient 0.4, p-value 0.2.

K-562 vs HL-60: Log2FC correlation coefficient 0.97, p-value 3.45 x10-8.

K-562 vs HL-60: Log2AS correlation coefficient 0.92, p-value 7.73 x10-6.

CHRF vs HL-60: Log2FC correlation coefficient 0.7, p-value 0.19.

CHRF vs HL-60: Log2AS correlation coefficient 0.5, p-value 0.39.

### MPRA lasso regression on annotated features

Enformer values^28^ were obtained from the vcf file of the 94 variants screened in the MPRA and cell matched features were selected for K-562 cells (455 features) and HL-60 cells (6 features). GWAS parameters, orthogonal scores and cell-matched Enformer features were combined in a unique matrix per variant to perform lasso regression on the continuous variable log2FC. We performed the lasso regression for ten iterations and selected predictive variables that had coefficients greater than 0 in at least three. Positive coefficients indicate that higher values of the sequence feature are predictive of higher values of Log2FC. Conversely, negative coefficients indicate that higher values of the sequence feature are predictive of lower values of Log2FC.

In the case of the lasso regression for predictive variables of a qualitative variable, first we transformed the labels into integers (MPRA negative RNA+ = 0 and Double positive = 1) and then we ran the lasso regression with all the features and selected those that had coefficients greater than 0 in at least one iteration. Positive coefficients indicated that higher values of the sequence feature are predictive of the double-positive class. Conversely, negative coefficients indicate that higher values of the sequence feature are predictive of belonging to the MPRA- /RNA+ class. The expression values for the K-562 genes in the basal state were obtained from ENCODE^41^.

### INTERVAL Differential Expression linear model

We modelled FPKM normalised raw values for each gene with a linear model using as covariates the PEER factors (top 35 from 50), top 10 genotype principal components, sex, age, BMI, RIN, sequencing batch, RNA concentration, read depth, and season (based on month of blood draw). Depending on the blood phenotypes associated with the variants at PP >= 0.1 we included a set of cell-count specific covariates (see **Table S1**) to account for cell count effects on total blood composition.

For each variant, we tested a median of 12 genes by combining all expressed genes within the GWAS association blocks (median size of 0.5 Mb) and those connected with index variants through PCHi-C interactions^45^. The multiple testing correction was done using the Benjamini– Hochberg method at three levels for each variant: i) all transcripts of all the genes in which the variant had a direct VEP Most Severe Consequence (LOF, MISS, SYN, UTR5, UTR3, INTRON, SPLICE, UPSTREAM), ii) all transcripts of all the genes within the GWAS association block plus the genes connected to the variant via PCHi-C (Block and PCHiC levels) and iii) Only for variants in Table 1, all the transcripts of all the genes (genome wide).

### INTERVAL Alternative Transcript Usage (ATU) additive logratio model

We calculated the median transcript ratio (median transcript TPM/median Expression of the gene to which the transcript belongs) for homozygous reference and heterozygous carriers separately. We discarded all transcripts with median transcript ratios below 0.1 in both genotypes to filter out transcripts whose contribution to the total expression of a given gene remains low irrespective of the genotype.

Given the compositional nature of the data we decided to transform proportions using the additive logratio model. Briefly, for all of the transcripts belonging to the same gene we first estimated the transcript ratio of gene expression (expression of the transcript/sum of the expression of all the transcripts belonging to the same gene) and scaled it to a reference transcript. The most abundant transcript was chosen as the reference transcript for each gene to avoid having 0 values at the denominator. This proportion was then log transformed. To avoid having transcripts with 0 TPM value in the logratio model, for any given transcript we imputed the value of the samples with 0 TPM to 0.65 of the minimum value greater than 0 for that transcript as this value is suggested to limit the distortion of the covariance matrix^87^. The resulting log scaled ratio per transcript was used in a linear regression model with the same constitutive and cell count specific covariates (see **Table S1**) per variant as the ones used in the DE model.

The multiple testing correction of the ATU model was done using the Benjamini–Hochberg method at two levels for each variant: i) all transcripts of all the genes in which the variant had a direct VEP Most Severe Consequence (LOF, MISS, SYN, UTR5, UTR3, INTRON, SPLICE, UPSTREAM), ii) all transcripts of all the genes within the GWAS association block plus the genes connected to the variant via PCHi-C (Block and PCHi-C levels). To represent the transcripts we used ggtranscript^88^.

### BluePrint Differential Expression linear model

We used the normalised gene quantification values from BLUEPRINT data^3^. We then applied a linear model to the gene quantification values for each gene.

The multiple testing correction was done using the Benjamini–Hochberg method at two levels for each variant: i) all transcripts of all the genes in which the variant had a direct VEP Most Severe Consequence (LOF, MISS, SYN, UTR5, UTR3, INTRON, SPLICE, UPSTREAM) and ii) all transcripts of all the genes within the GWAS association block plus the genes connected to the variant via PCHi-C (Block and PCHiC levels).

### BluePrint Alternative Transcript Usage (ATU) additive logratio model

We calculated the median transcript ratio (median transcript FPKM/median Expression of the gene to which the transcript belongs) for homozygous reference and heterozygous carriers per cell type separately. For every cell type of BLUEPRINT we discarded all transcripts with median transcript ratios below 0.1 in both genotypes.

The same compositional model and multiple testing correction used for whole blood was applied in the BLUEPRINT per cell type.

### INTERVAL GSEA and ORA

For the GSEA analysis we started from the 22 variants in **Table 1** and performed a genome wide DE analysis between carriers and non-carriers per variant in the INTERVAL whole blood dataset. Next, we used the Fold Change between WT and HET genotypes to order the genes in a decreasing manner and input the ordered gene list into the GSEA function of ClusterProfiler^89^. The minimum and maximum gene set size and the p-value cutoff were set to 10, 500 and 0.05, respectively. The multiple testing was accounted for by Benjamini & Hochberg.

For the ORA analysis we started by defining a list of gene sets for blood and immunity from the Molecular Signatures Database (MSigDB)^90^: we first selected all the gene sets that contained in their description at least one of the following blood and immunity related terms: PLATELET, ERYTHRO, MEGAKARYOCYTE, MONOCYTE, NEUTROPHIL, EOSINOPHIL, BASOPHIL, LYMPHOCYTE, T_HELPER, TH_17, TH17, TH1, TH2, BLOOD, BLOOD_COAGULATION, IMMUNE, HUMORAL_IMMUNE_RESPONSE, IMMUNOGLOBULIN and HEMATOPOIETIC. We also included two blood and immunity unrelated terms (HEPATOCYTE and NEURON). In addition, we included in the ORA analysis TF target gene sets for the TFs *GATA2, GFI1, CUX1* and *RUNX1* in the Dorothea collection^91^ (A,B,C and D confidence levels). The minimum and maximum gene set size allowed was 10 and 500, respectively, but for the Dorothea gene sets which were exempted from this filter (*CUX1* n=464 genes, *FOXP1* n=3,278, *GATA2* n=5,370, *GFI1* n=8 and *RUNX1* n=2,551). Next, we tested the genome wide DE analysis of the 22 variants in **Table 1** used for the GSEA for overrepresentation of DE genes applying Active Pathways^92^ with the significant p-value cutoff set to 0.01. As a background list of gene sets we used all the gene sets derived from the Human Phenotype ontology (c5.hpo.v2023.2.Hs.entrez.gmt, *n=5,547 gene sets*^36,90^). We obtained 64 pathways significantly overrepresented in DE genes of which 3 were from the unrelated terms. The remaining 65 pathways corresponded to 11 variants. We excluded from further analysis the gene sets from Dorothea (2 variants). A variant was labelled as ‘ORA positive’ if it had at least 1 ORA pathway at p-value = 0.05. To test if the 9 variants were enriched in the TF annotation (**Table 1**) we used a Chi square test (6/7 variants with TF annotation vs 3/15 non TF labelled variants were ‘ORA positive’, p-value = 0.0141).

### DE and ATU analysis on bulk RNA-seq of clone lines carrying the reference (G/G) and homozygous alternative (C/C) genotypes for rs35592432

Adapter trimmed reads were aligned against the human genome (GRCh38 Ensemble release 115, annotated with gencode v49 gtf) using the splice aware aligner STAR^93^. Gene and isoform level quantifications were obtained using RSEM^94^. To model ATU we used DRIMSeq^30^ given the absence of covariates due to the simple experimental design. To calculate DE we used DESeq2^95^ and for ORA we employed the enricher function from the package clusterProfiler^89^. We selected as gene sets all the Dorothea^91^ *FOXP1* target collections that comprised between 10 and 500 genes.

### Comparison of number of carriers between variants with DE/ATU regulation and variants without effect in the RNA-seq studies

Median number of heterozygous carriers for variants with DE and/or ATU = 43 and median number of heterozygous carriers for variants without RNA-seq effect = 36, p-value = 0.045, *Wilcoxon test*.

### GenIE analysis

Read pairs per amplicon were merged using flash^53^ with the following parameters: read length (-r) 150, fragment sd (-s) 20, minimum overlap (-m) 10. Fragment size (-f) and maximum mismatch density (-x) depended on the targeted amplification: *CUX1* (223, 0.115) and ENSG00000178927/CYBC1/EROS (249, 0.12). The reads were aligned using bwa mem^96^ with the following parameters: -O 24,48 -E 1 -A 4 -B 16 -T 70 -k 19 -w 200 -d 600 -L 20 -U 40. The aligned reads were filtered to discard reads with more than 10 clipped bases using samclip^97^. The GenIE results were obtained using the rgenie package^12,98^ allowing for 10 bases for the required_match_right and required_match_left parameters.

### Flow cytometry data ILR compositional analysis

We collected the percentages of CD41^-^CD235^-^, CD41^-^CD235^+^, CD41^+^CD235^+^, CD41^+^CD235^-^ cells and the value of CD41 MFI and GeoMFI at the four time points of the PMA differentiations per genotype and clone line (*n=3* clone lines per genotype). The ‘Basal’ time point was assigned to cells mock treated for 16 hours. We used the package Compositions^99,100^ to transform the cell abundance percentages into isometric log ratios (ILR)^101^. We modelled the ILR values using either time (reduced model) or time and genotype (full model). We then tested by ANOVA if the genotype significantly improved the fit of the model for each pairwise comparison between WT genotype and the three edited genotypes. The p-values for the different models are as follows:

G/G vs A/A: Anova p-value = 1.54e-08

G/G vs ‘80 bp del’: Anova p-value = 6.98e-04

A/A vs ‘80 bp del’, Anova p-value = 2.04e-06

### Cell size using flow cytometry FSC-A and SSC-A

We collected the values of FSC-A and SSC-A per cell, genotype and time point and restricted our analysis to CD41^+^CD235^-^ cells at 72 hours as they were the most mature cell type in our model (*n=4,892, 8,914* and *7,553* for the wild type clone lines, *n=3,616, 4,307* and *6,229* for the homozygous alternative clone lines and *n=9,822, 6,476* and *9,611* for the 80 bp deletion clone lines). The median values for FSC-A are: 585312 (‘G/G’ genotype), 592815 (‘A/A’ genotype) and 647251 (Del80 genotype). The median values for SSC-A are: 391532 (‘G/G’ genotype), 407041 (‘A/A’ genotype) and 468646 (Del80 genotype). We used the Wilcoxon-rank test to analyse differences between the cells.

### 10X Multiome DE and DA analysis

Briefly, for the DE analysis we aggregated the counts of all the cells belonging to the same combination of clone line, time point and Seurat cluster using the function Seurat2PB (Seurat v5.01^75^). Then, per Seurat cluster, we used DESeq2^95^ to calculate a linear model accounting for time (reduced model) or for time and genotype (complete model) and applied a Likelihood ratio test (LRT) to test cases in which the complete model was a significantly better fit to the data following analysis guidelines for time and condition differential analysis^102,103^.

For the ORA analysis we started by defining a list of gene sets relevant for K-562 differentiation from the Molecular Signatures Database (MSigDB)^90^: we selected all the gene sets that contained in their description at least one of the following terms: PLATELET, ERYTHROCYTE, CUX1, MEGAKARYOCYTE, GATA1, GATA2, TET2, RUNX1, RUNX2, MITOSIS, ANEUPLOIDY, CYTOKINESIS, MYELOID, AML, LIPID, SPHINGOSINE, FOXM1, SPI1, PU1, PI3K, AKT, FOXP1 and GFI1. In addition, we included in the ORA analysis TF target gene sets for the TFs *GATA2, GFI1, CUX1* and *RUNX1* in the Dorothea collection^91^ (A,B,C and D confidence levels). The minimum and maximum gene set size allowed was 10 and 500, respectively.

For the DA analysis we first extracted all the linked peaks to the set of DE genes and a set of marker genes (in total 1,897 genes) using the LinkPeaks function of Signac (v1.12.0)^76^. To characterise the linked peaks (n=14,211 peaks) we overlapped them with annotated gene TSS (+/- 2.5 kB of the TSS, n= 4,072 overlaps with a known TSS, ENSEMBL, release 111^104^) and with the annotated regulatory features of the basal state K-562 cells (ENSEMBL, release 111^105^). Next, we produced a reduced seurat object with the ATAC counts of the selected peaks using the function CreateChromatinAssay (Signac v1.12.0) and then obtained a new Seurat object with the assay option set to ‘RNA’ as the function Seurat2PB (Seurat v5.01) would not work if it was set to ‘ATAC’. From this point onwards the pipeline proceeded as explained in the DE analysis. To clusterize the results of both analyses we used Pheatmap^106^ and to display them in volcano plots and in detailed locus plots we used ggplot2^107^. The results of the DE and DA analyses are included in the Zenodo repository.

The chromVAR^38^ analysis was carried out using the RunChromVAR function of Signac and then subsetting all the cells in clusters 1 and 3 and running FindMarkers function for every genotype against the wt using the t test. To obtain gene regulatory networks (GRNs) for *CUX1* and *RUNX1* in the multiome dataset we pre-processed the data using SIMBA^108^ and then identified GRNs using their guidelines.

## Additional resources

<u>Code to recreate the computational environments, reproduce the analyses, access to raw and intermediate data, and reproduce all figures and tables</u>

https://zenodo.org/records/16539111

<u>MPRA library synthesis and cloning protocol:</u> https://www.protocols.io/edit/mpra-synthesis-library-design-and-cloning-soranzo-cs3awgie

Step by step protocol used to design and clone the MPRA oligos into the reporter vector.

<u>MPRA nucleofection:</u> https://www.protocols.io/edit/mpra-synthesis-cellular-work-and-nucleofection-sor-cs3jwgkn

Step by step protocol used to nucleofect the library into the cancer cell lines.

<u>MPRA library preparation for sequencing</u>: https://www.protocols.io/edit/mpra-synthesis-dna-rna-isolation-and-library-prepa-cs3mwgk6

Step by step protocol used to extract and prepare the different MPRA libraries for sequencing.

<u>Haemvar architecture/ documentation:</u> https://haemvar.org

We created the HaemVar Database of genetic variants and blood-related traits, containing the complete association and fine-mapping result data set of all the variants associations to 29 blood cell phenotypes assessed by Vuckovic et al^2^, and including further annotation data for the prioritised subset of 178,890 variants as described above; see section ‘Variant Annotation’. All information contained in the HaemVar Database is presented through an open-access website that generates comprehensive gene and variant-specific data reports with downloadable tables and figures. The software of the web-based application is written in PHP for server-side handling and optimisation of database access and data output, and uses the open-source Vega v5 (http://vega.github.io) JavaScript library for custom generation of data-driven and (partially) interactive visualisations.

Declaration of generative AI and AI-assisted technologies in the writing process

During the preparation of this work the author(s) used ChatGPT in order to improve language and readability. After using this tool/service, the author(s) reviewed and edited the content as needed and take(s) full responsibility for the content of the publication.

