## Supplementary figures and images for "Integrating Natural and Engineered Genetic Variation to Decode Regulatory Influence on Blood Traits"

### Supplemental Figure 1

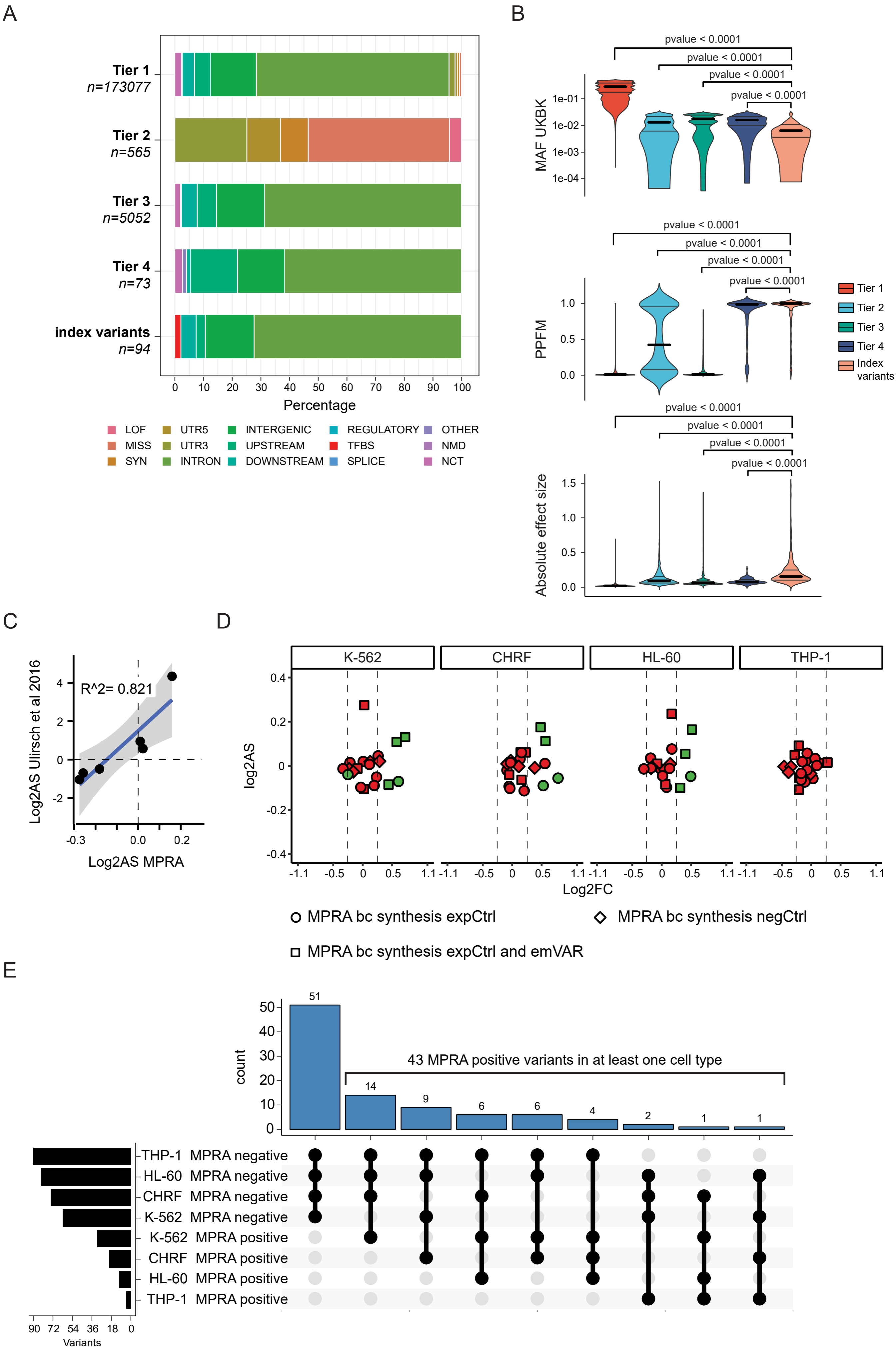

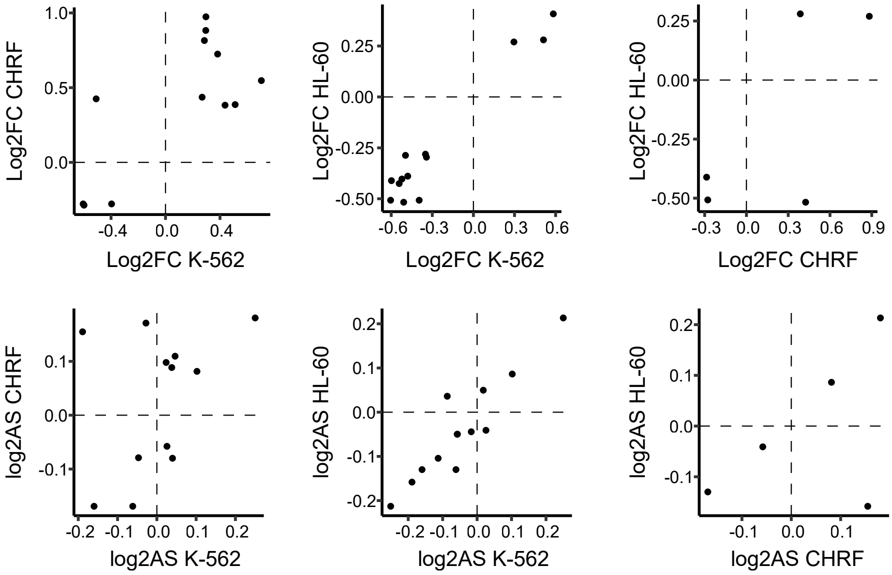

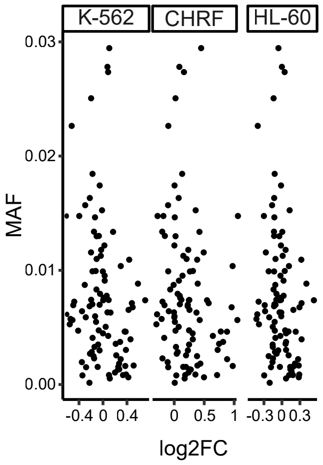

### Supplemental Figure 2

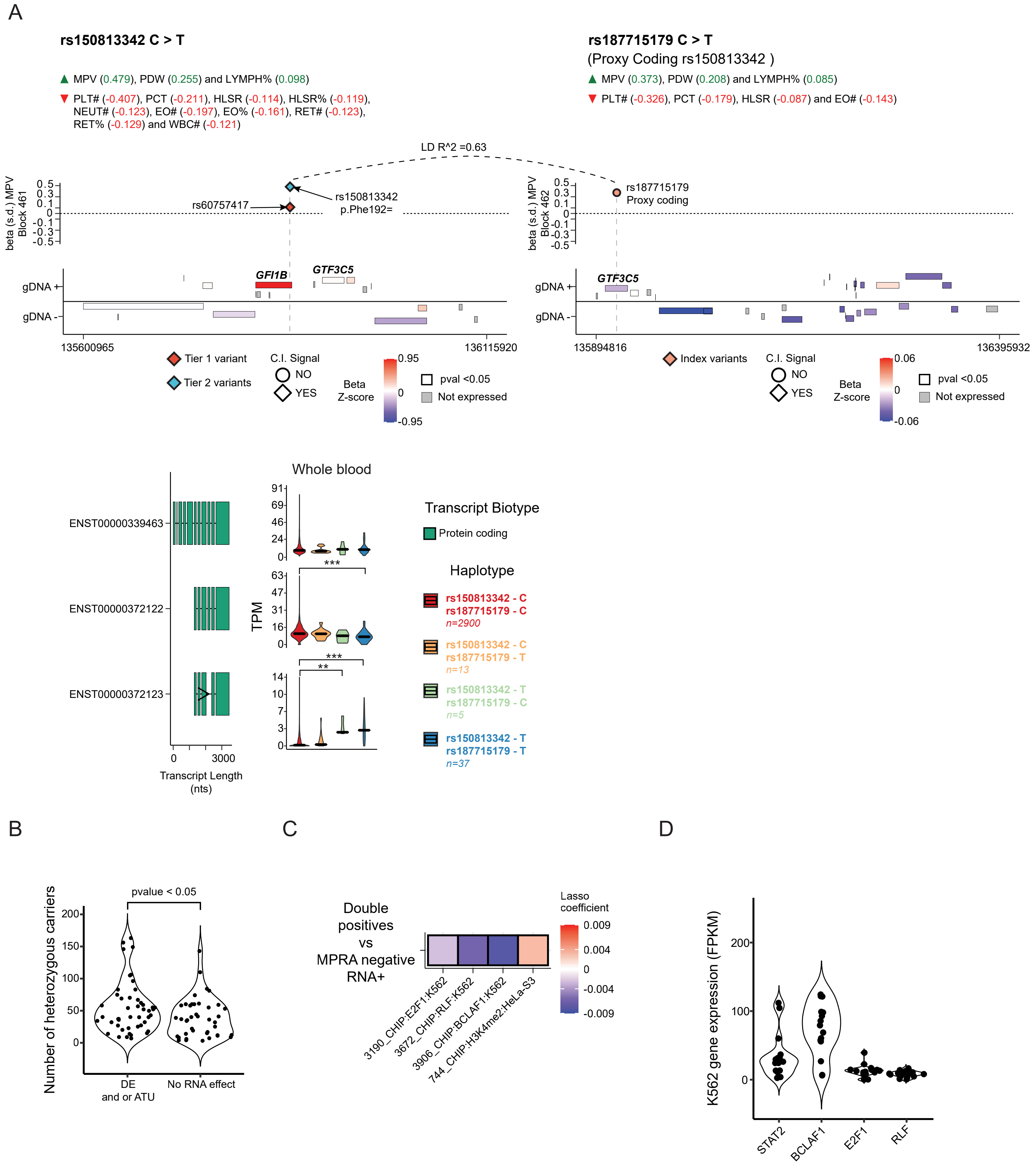

### Supplemental Figure 3

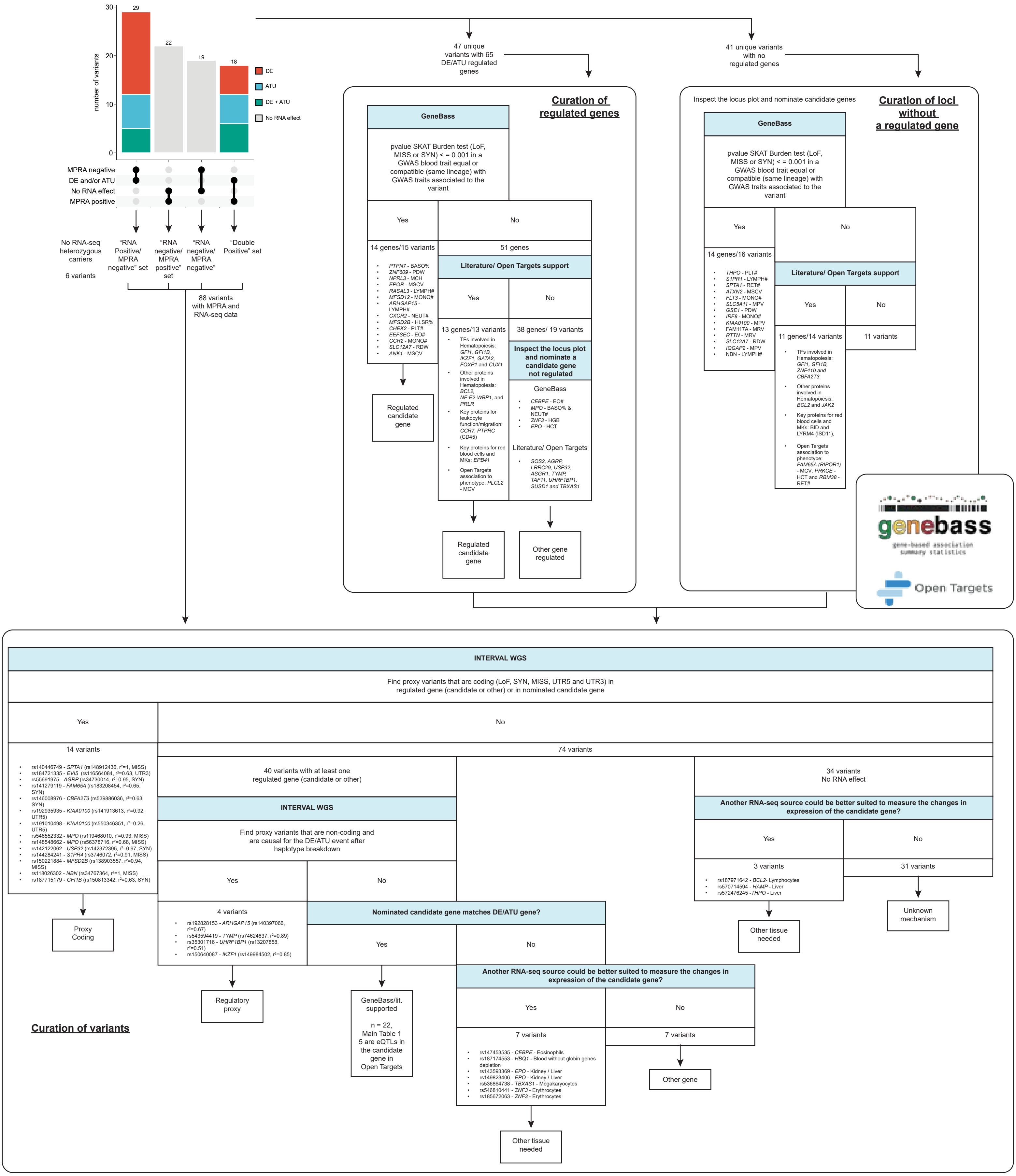

### Supplemental Figure 4

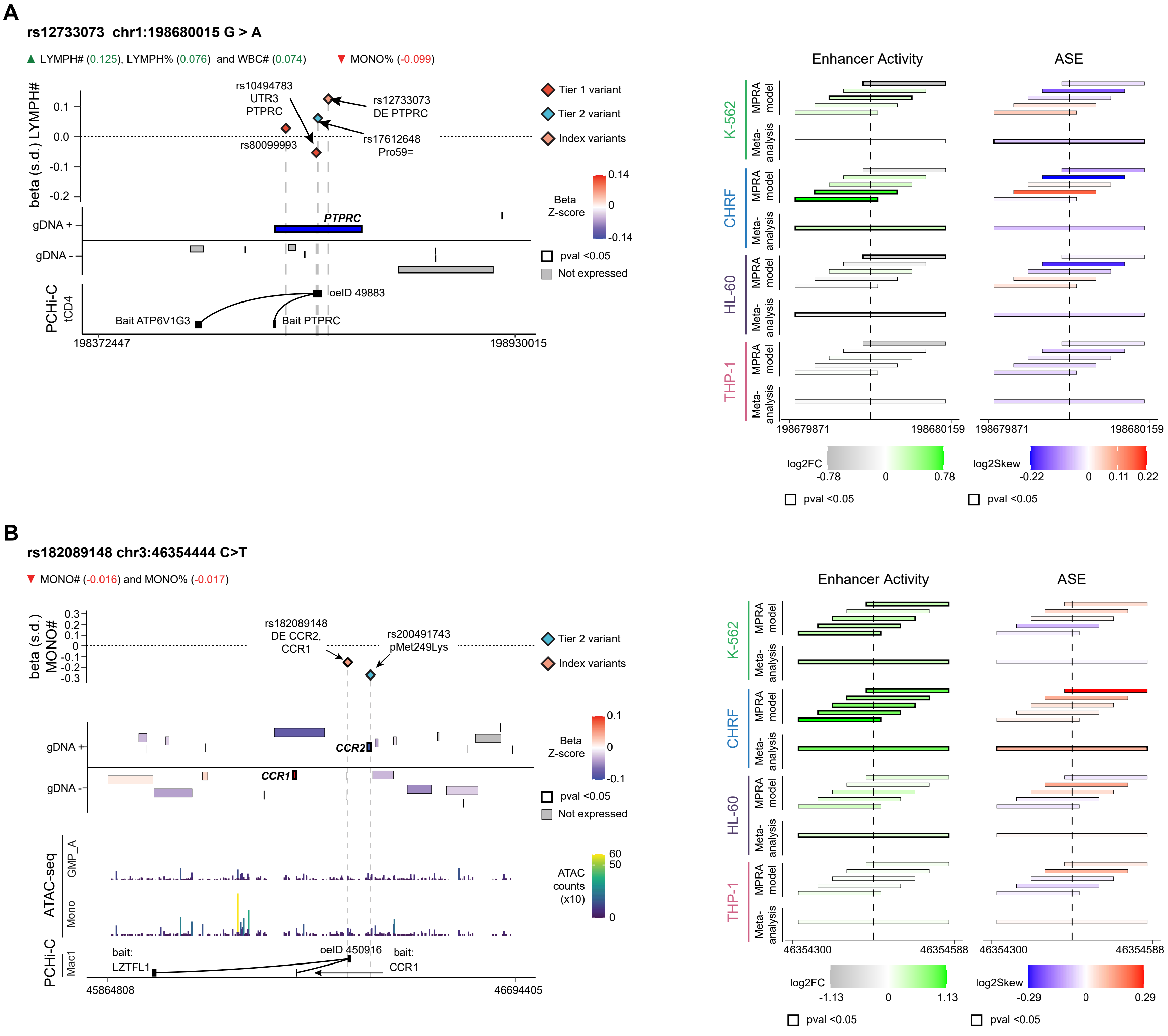

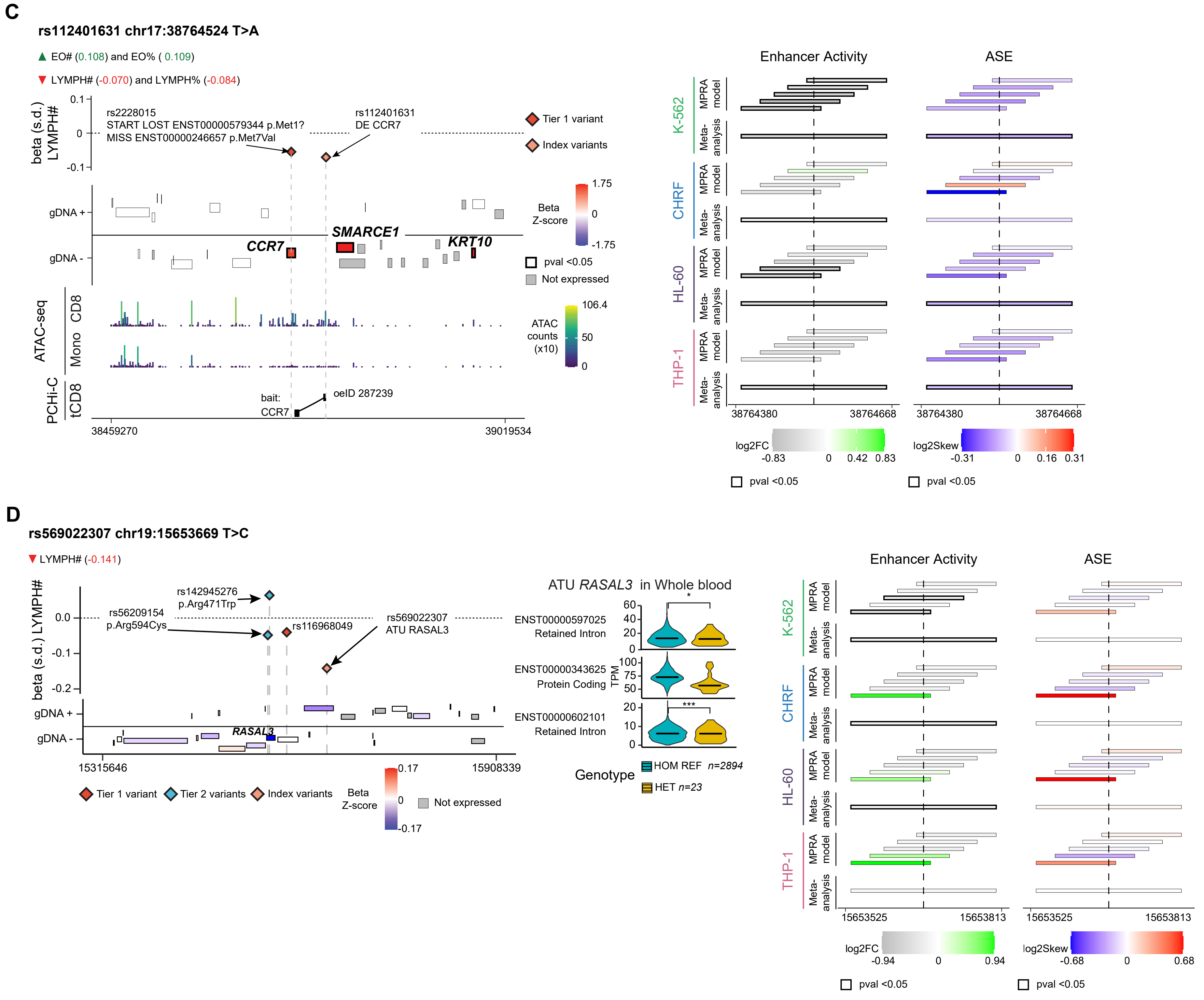

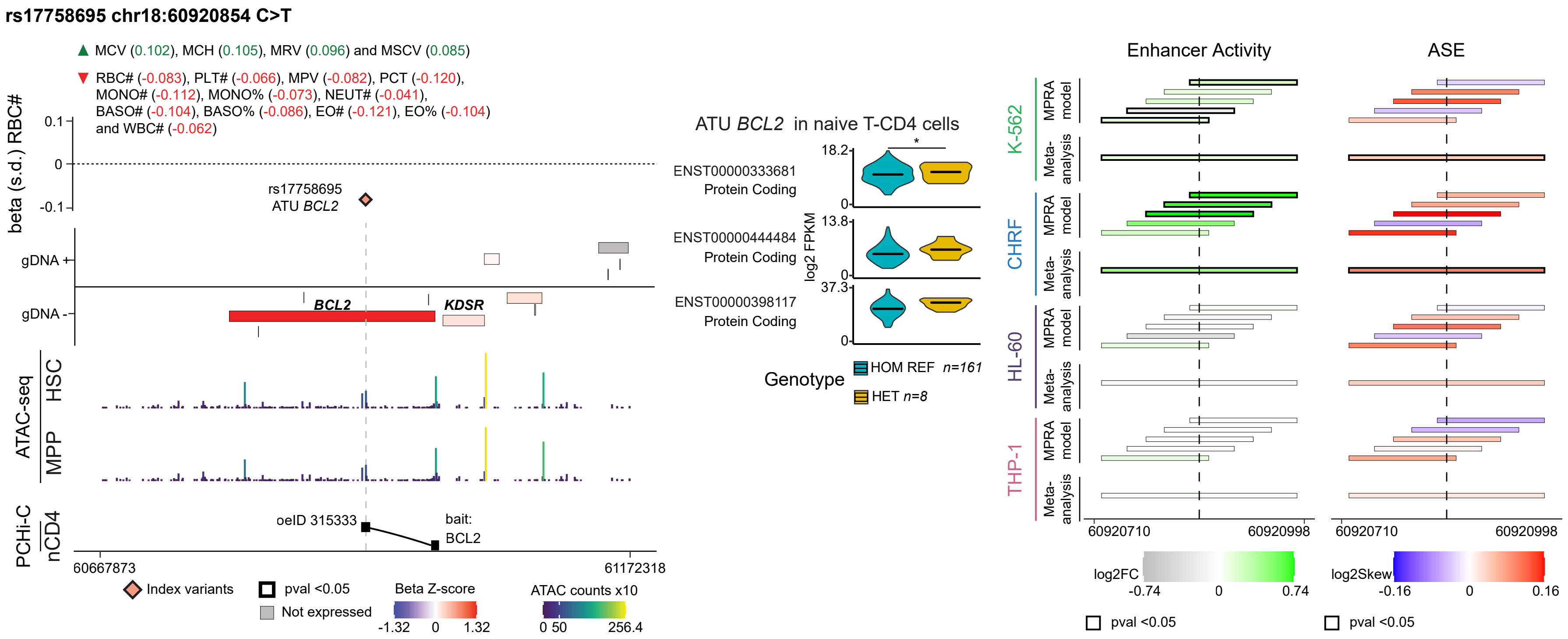

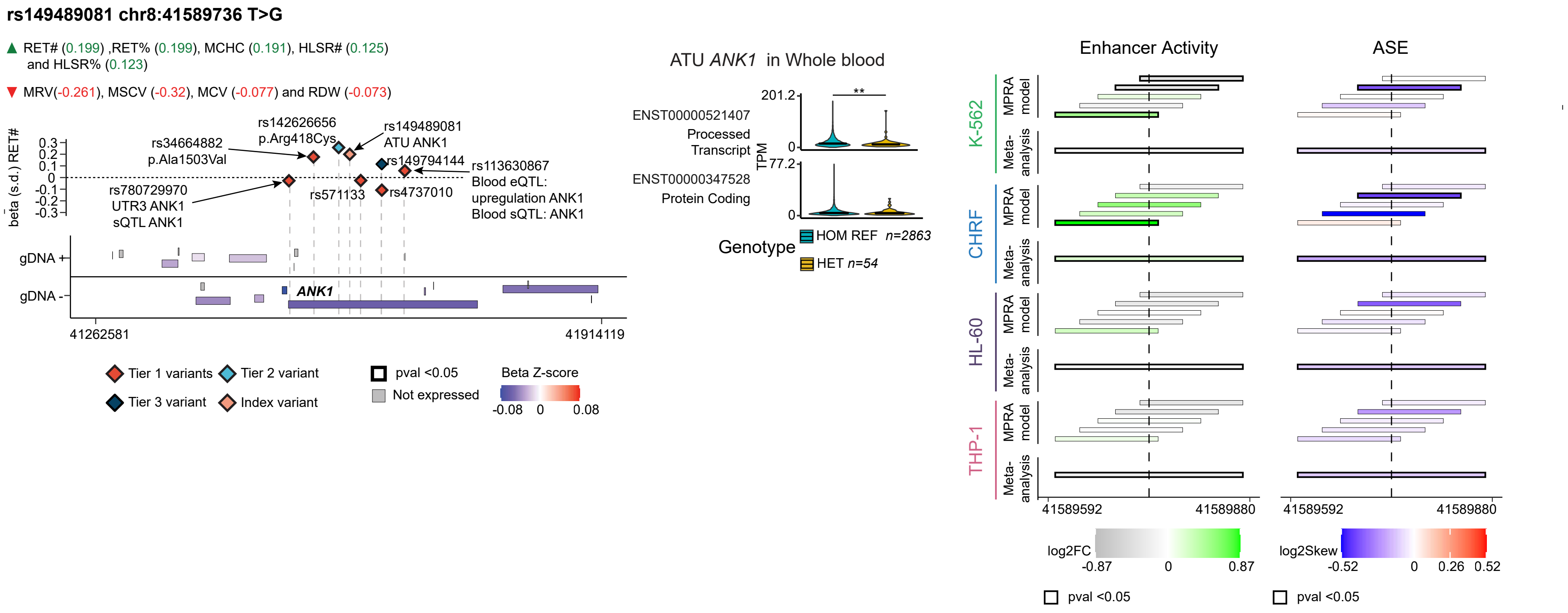

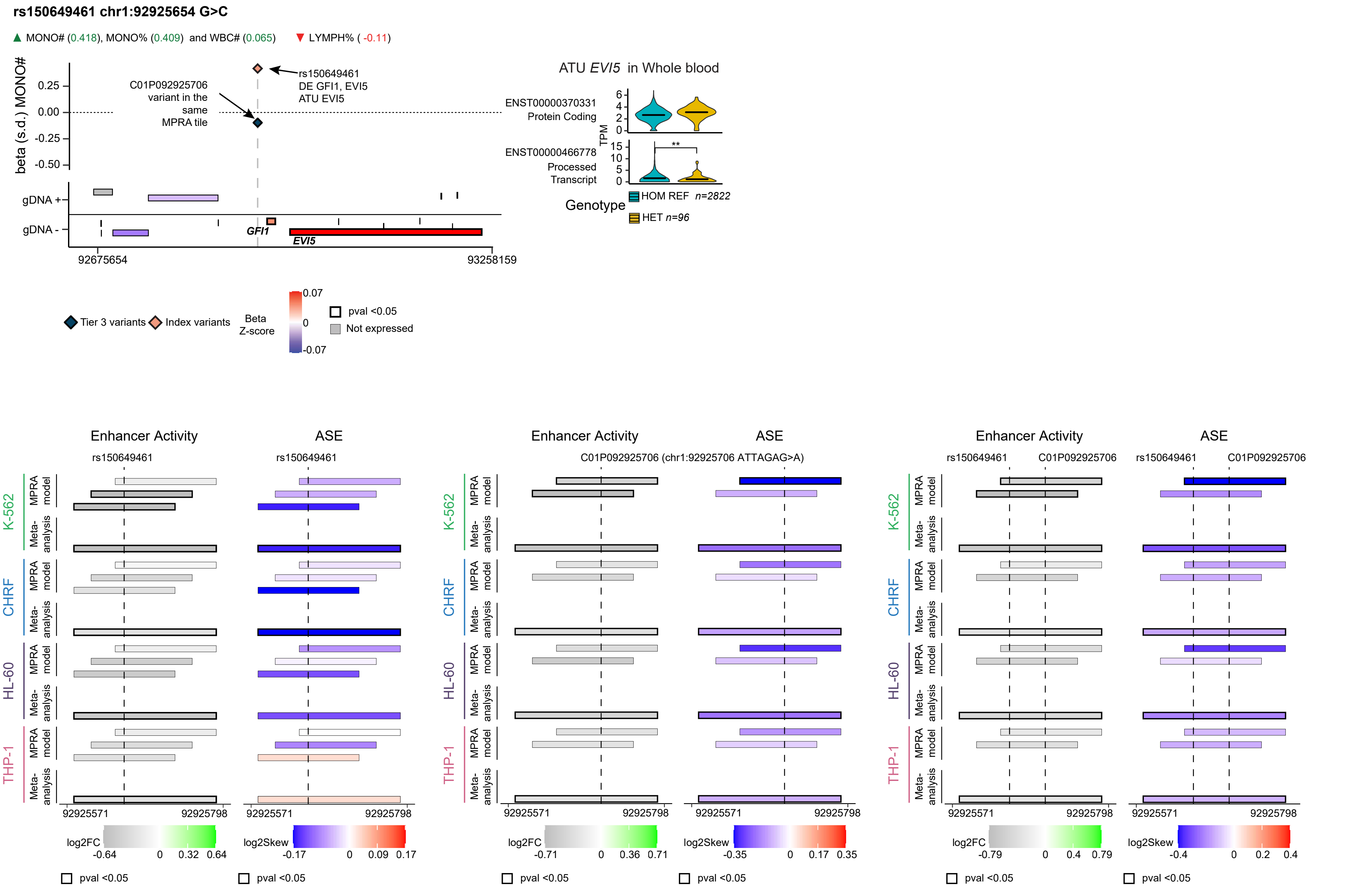

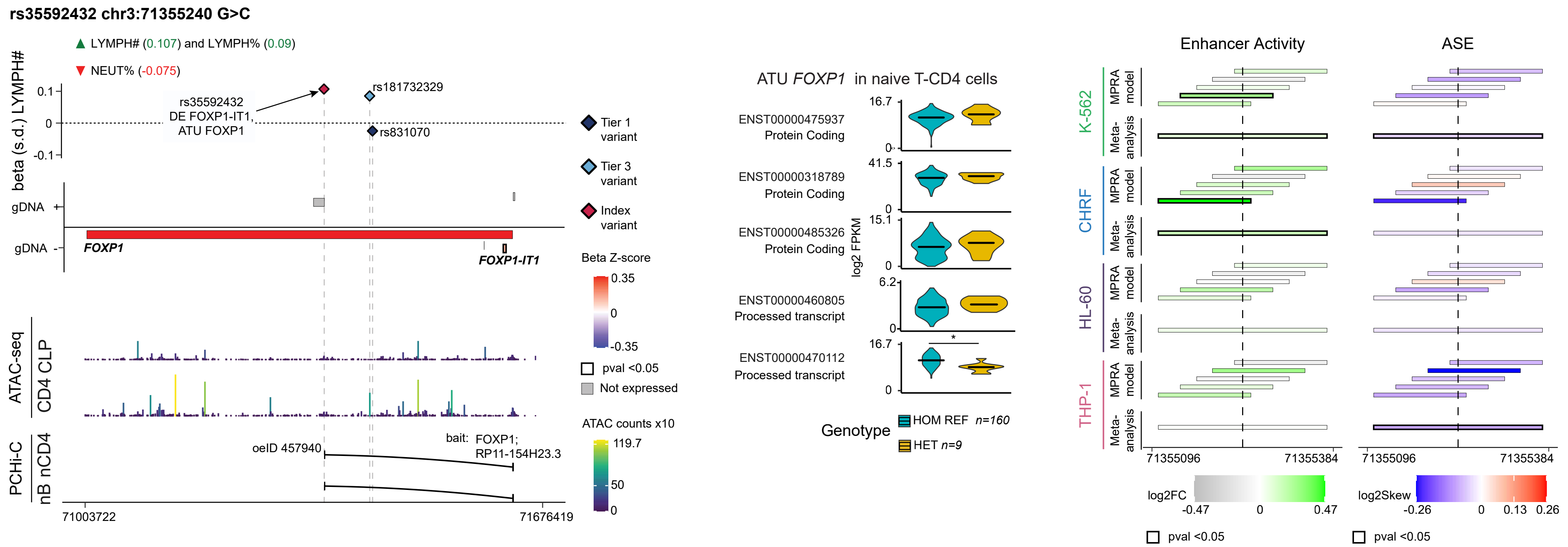

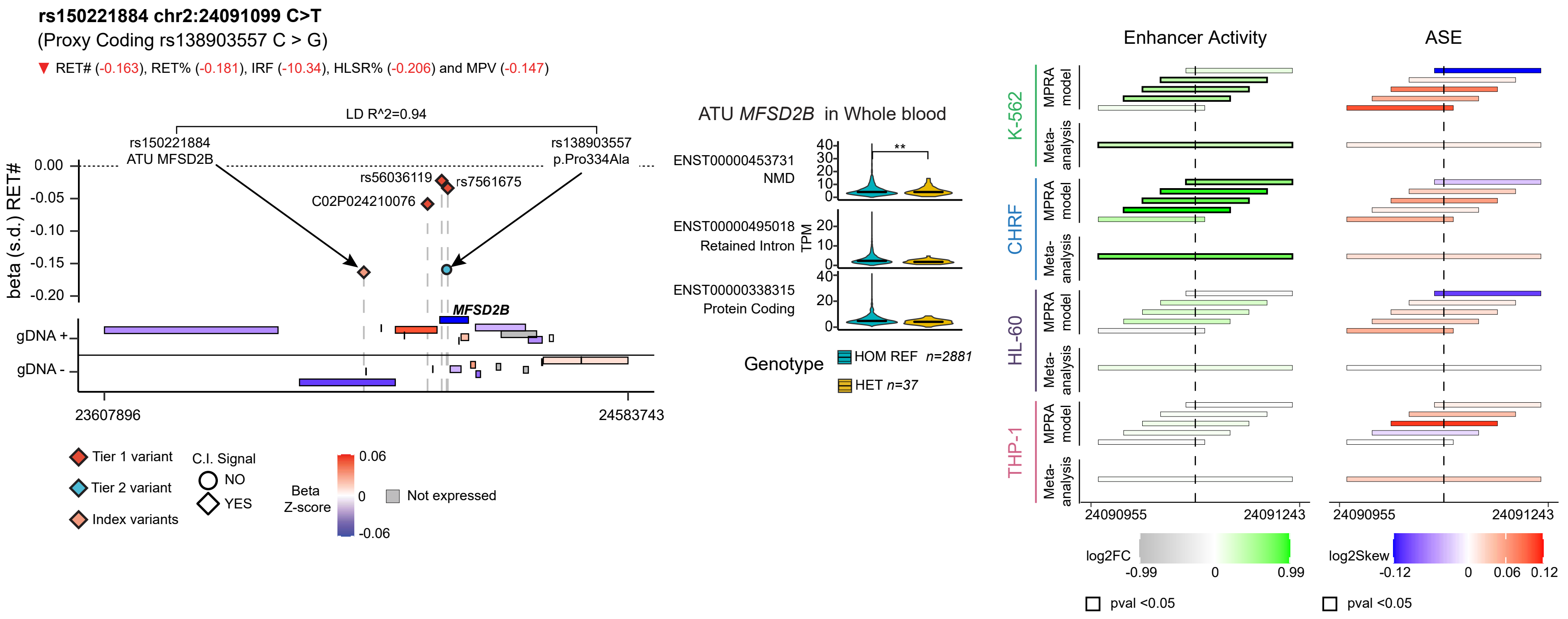

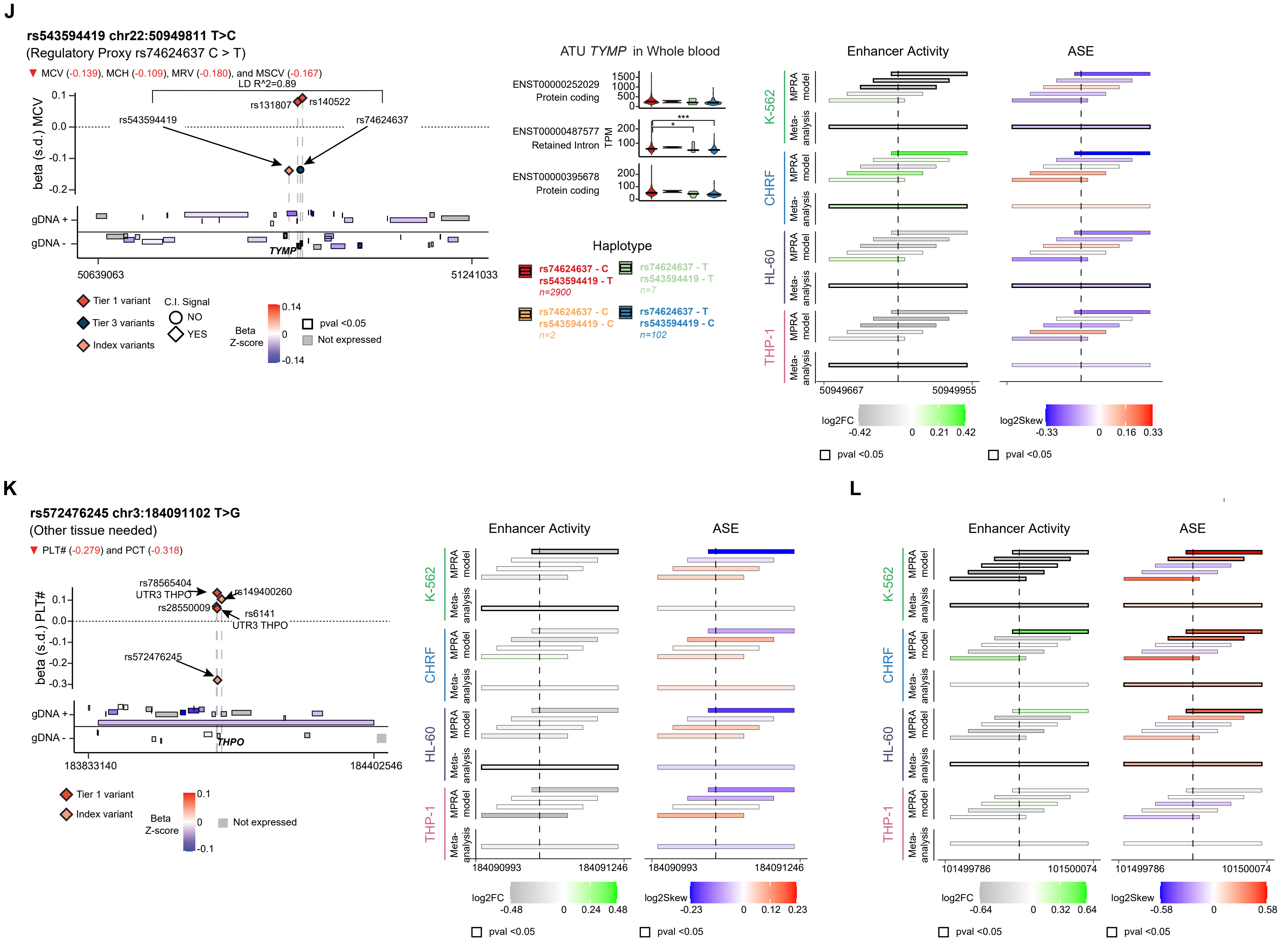

### Supplemental Figure 5

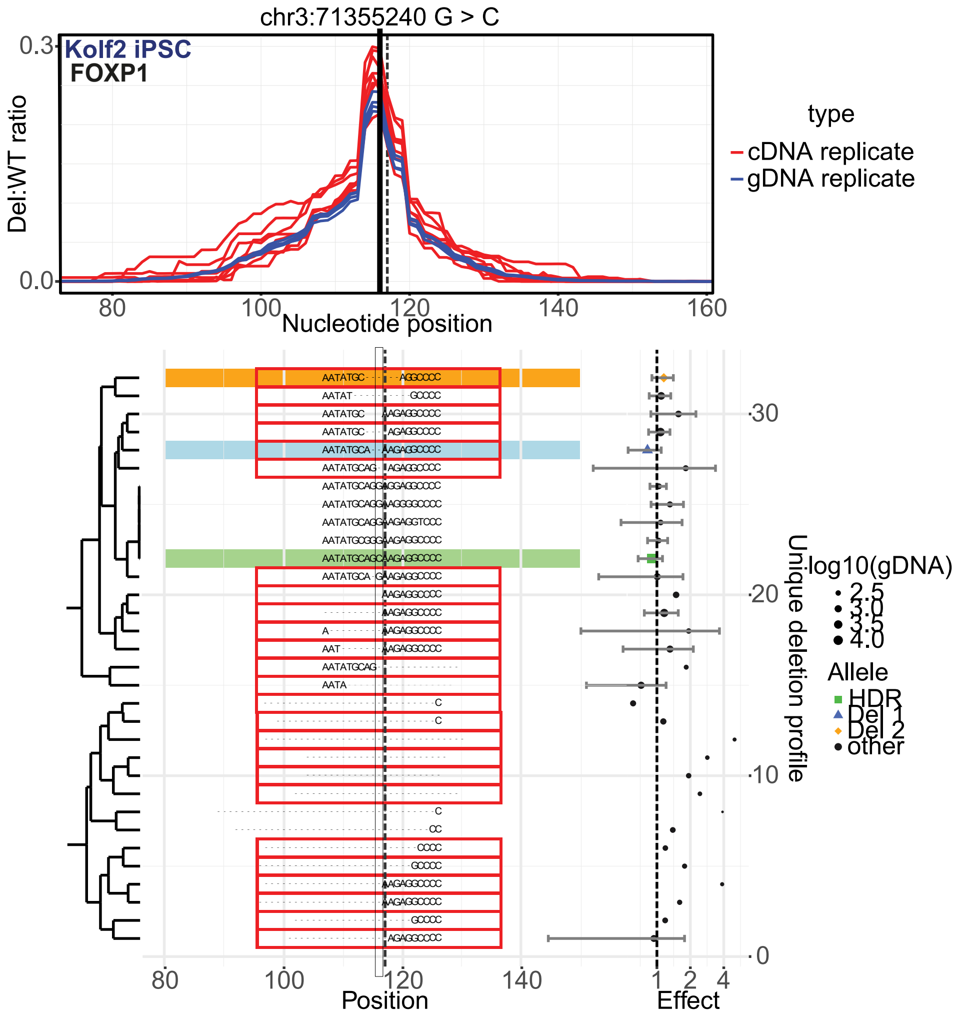

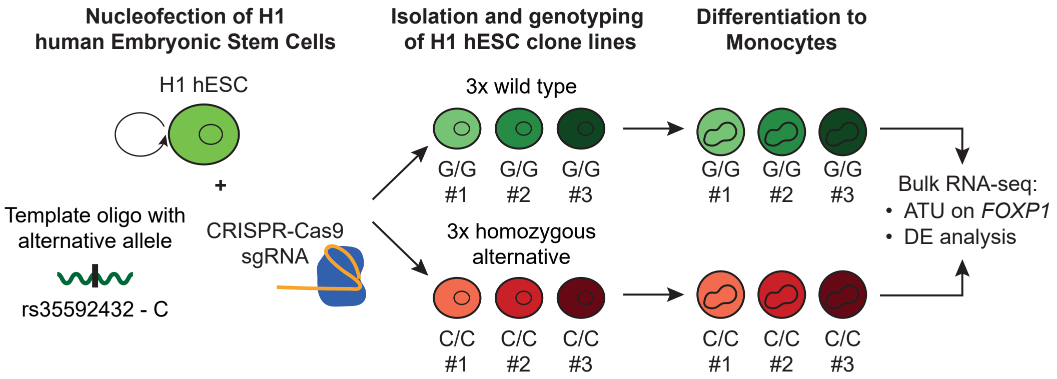

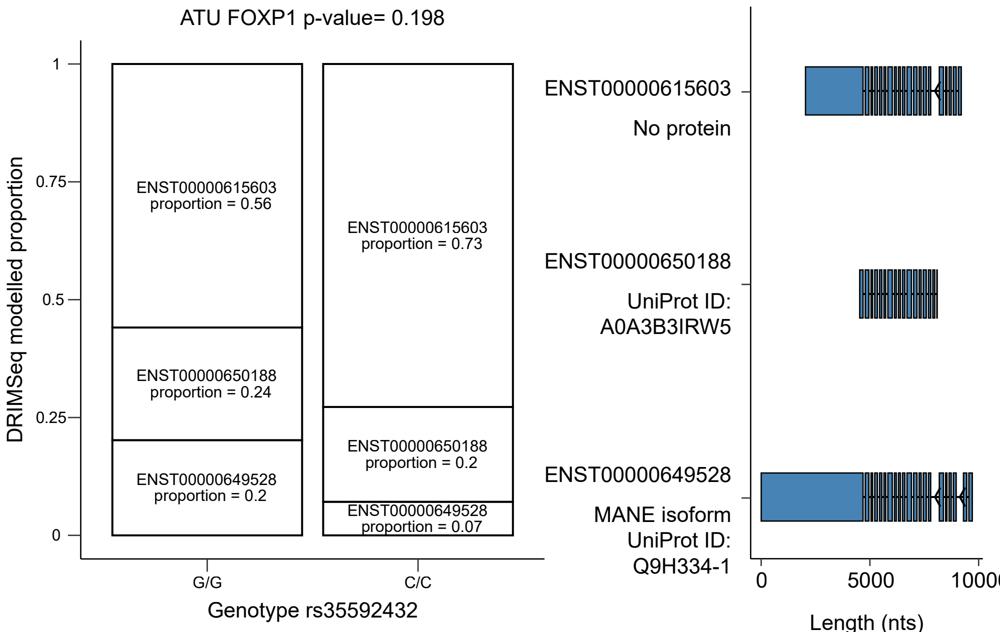

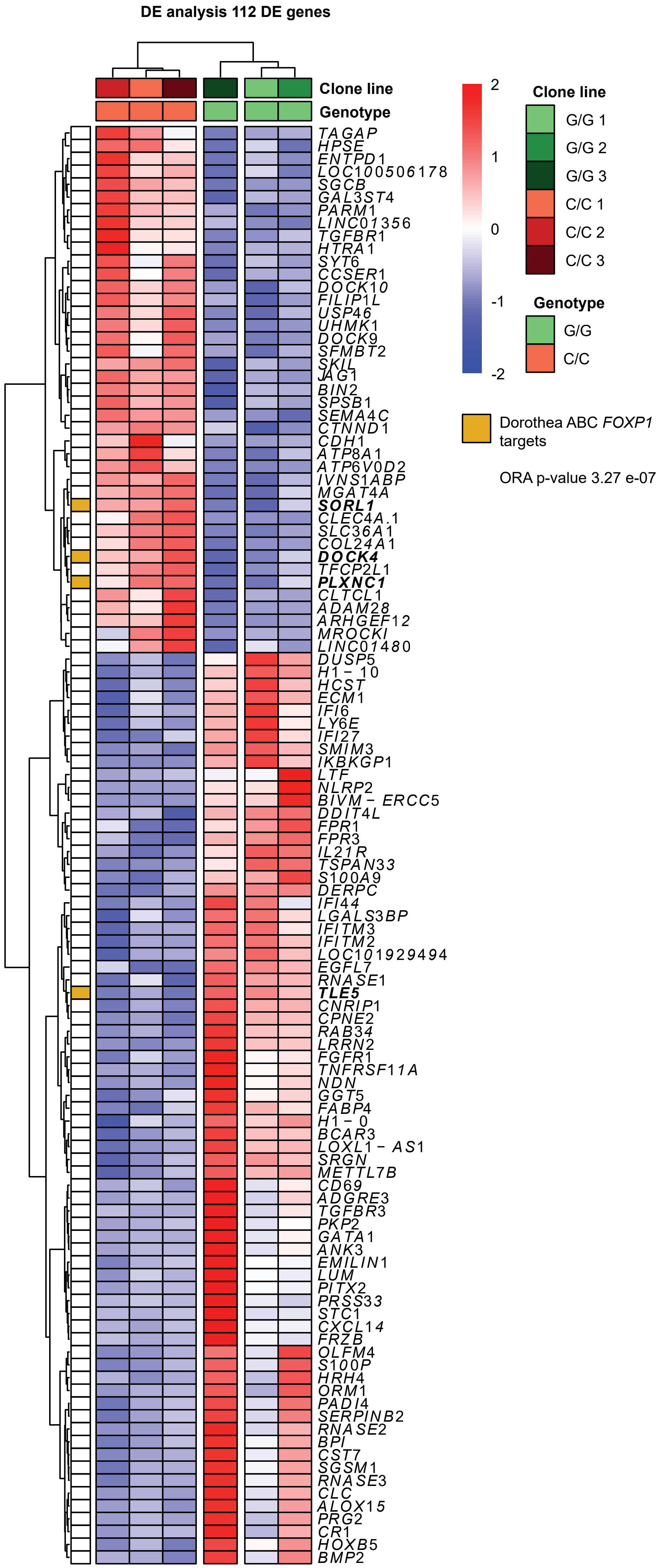

### Supplemental Figure 6

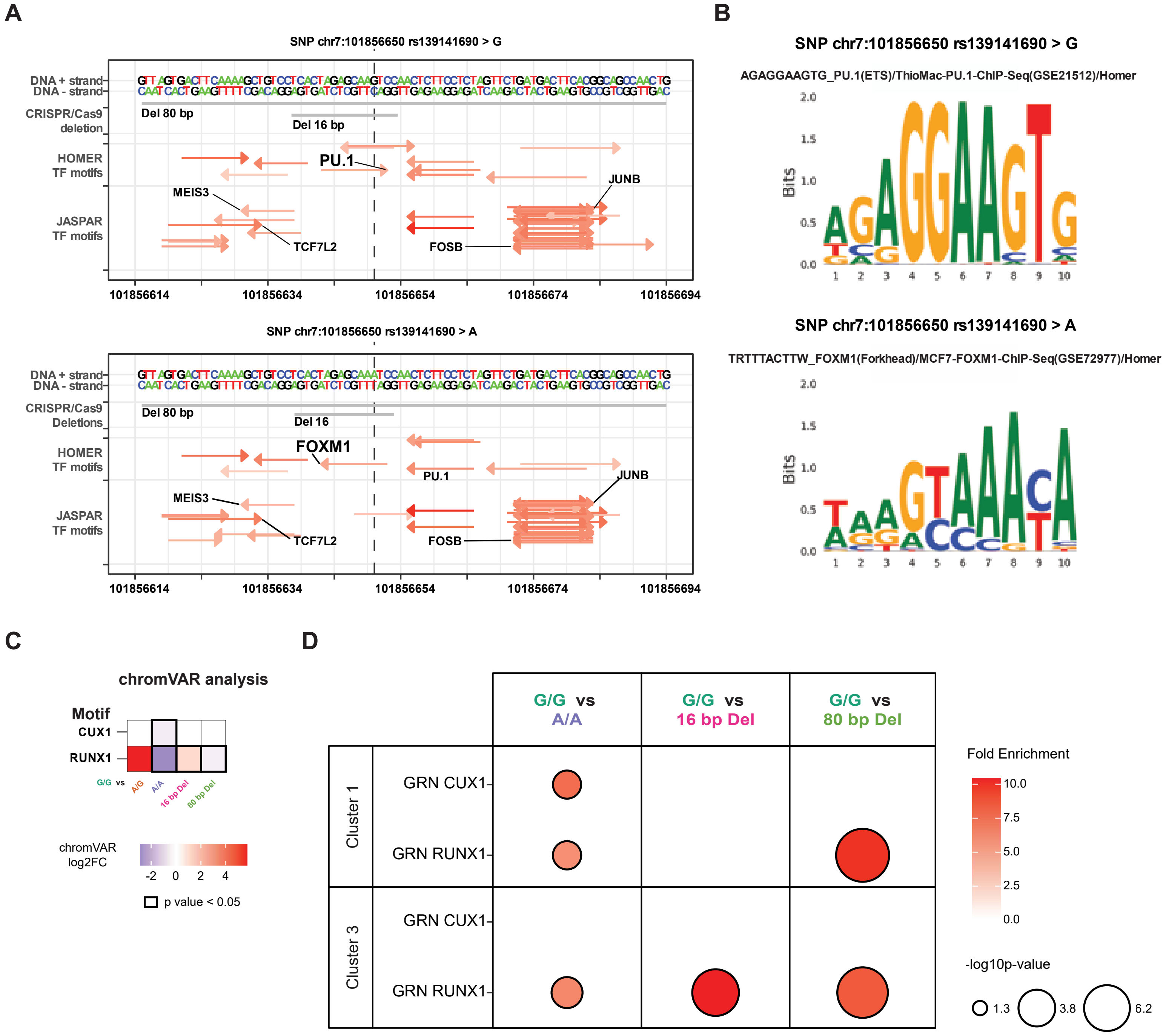

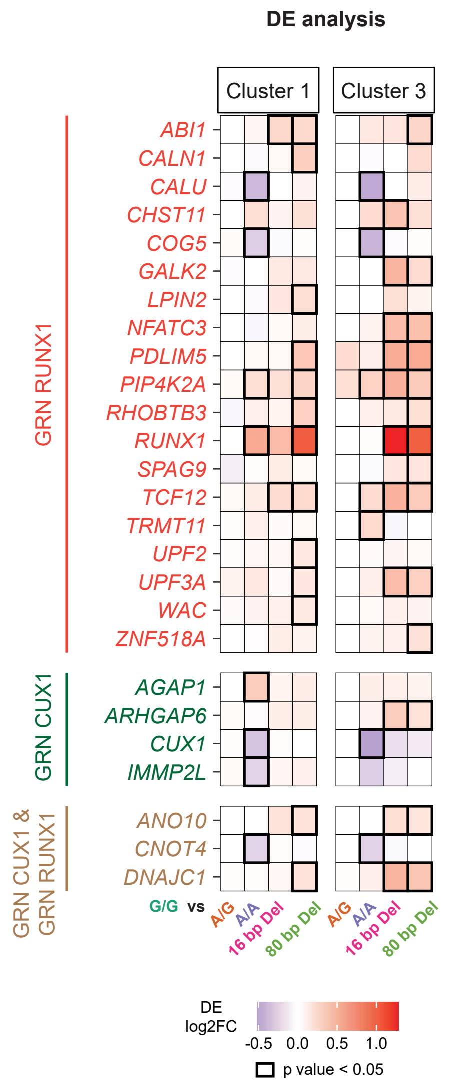
