## Supplemental Information for "Integrating Natural and Engineered Genetic Variation to Decode Regulatory Influence on Blood Traits"

**Supplementary Information**

**Supplementary figures**

**Supplementary Figure 1 related to Figure 2.** **MPRA results, selectivity and cellular specificity. A)** VEP most severe consequence for the variants in the different tiers. **B)** Values of the MAF from UK Biobank, PPFM and Absolute effect size across the different tiers. **C)** Comparison of ASE directionality in the set of overlapping variants between our MPRA in K-562 cells and a previous MPRA in the same cell type^1^. **D)** Adjustment of the log2FC threshold to maximize the amount of MPRA positive variants using MPRA positive controls from previous studies (MPRA bc synthesis expCtrl and MPRA bc synthesis expCtrl and emVAR^1^, **Table S3** and **S4**) while controlling for a set of four regions deprived of activity in CRISPR-A assays (MPRA bc synthesis negCtrl^2^, **Table S3** and **S4)**. The log2FC threshold was set at +/- 0.255. **E)** MPRA results after the meta-analysis of tiles broken down by MPRA positives and negatives per cell type. **F)** Pairwise correlation plots for log2FC and log2AS in K-562, CHRF and HL-60 cell lines across shared MPRA positive variants. THP-1 had too few shared MPRA positive variants. **G)** Values of MAF against log2FC for K-562, CHRF and HL-60 cell lines. Negative lasso regression coefficients for MAF indicated that lower values (MAF < 0.01) were predictive of higher log2FC. *MAF UKBK = minimum allele frequency from UK Biobank, PPFM = Posterior Probability in the Fine Mapping. The comparisons correspond to the values of the ‘index variants’ versus the rest of the tiers, Wilcoxon test. For the abbreviations of blood phenotypes see Table S3.*

**Supplementary Figure 2 related to Figure 3. Variants tagging coding and non-coding proxies that cause ATU.** **A)** Locus plots for the variants rs150813342 and rs187715179. rs187715179 tags the coding proxy rs150813342 which is a synonymous mutation in the *GFI1B* transcription factor. The ATU analysis revealed that the haplotypes rs150813342-T-rs187715179-C and rs150813342-T-rs187715179-T entail an ATU event that results in a lower contribution of a protein coding transcript (ENST00000372122, short protein) to the pool of transcripts of *GFI1B.* **B)** Variants without any regulation detected in the RNA-seq had significantly lower carriers in our datasets. **C)** Lasso regression on sequence features predictive of belonging to double-positive variants as opposed to MPRA-/RNA+. **D)** Expression of the transcription factors *STAT2, BCLAF, E2F1* and *RLF* in different samples of K-562 cells obtained from ENCODE^3,4^. Negative lasso regression coefficients indicated that low CHIP-seq values for *BCLAF, E2F1* and *RLF* were predictive of *double positive* variants whereas high CHIP-seq values were predictive of MPRA-/RNA+ variants. Both *E2F1* and *RLF* were lowly expressed in K-562 cells. *STAT2* is shown to provide comparison with the expression levels of a transcription factor whose CHIP-seq values were found to positively predict higher log2FC in the MPRA. *For the abbreviations of blood phenotypes see Table S1 and Table S2 for the breakdown of variant classification in tiers. * p value <= 0.05, ** p value <= 0.01, *** p value <= 0.001, ATU additive log ratio model.*

**Supplementary Figure 3 related to Figure 3. Flowchart of Manual curation.**

**Supplementary Figure 4 related to Figure 3. Manual curation excludes variants tagging coding and regulatory proxies.** **A)** Locus plot for the variants impacting the gene PTPRC (CD45) and the GWAS trait LYMPH#. Two of the variants (rs17612648 and rs10494783) are coding (CADD values 0.26-3.8) affecting the *PTPRC* gene. We observed a significant downregulation of *PTPRC* for the rs12733073-A allele. The meta-analysis of the MPRA shows that the variant is not MPRA active in any cell line. **B)** Locus plot for the variants impacting the genes *CCR1* and *CCR2* and the GWAS trait MONO#. The independent variant rs200491743 (Tier 2) is a coding variant affecting *CCR2*. ATAC-seq shows accessible chromatin in cell types belonging to the myeloid lineage whereas the PCHi-C track for the Mac1 cell type shows a region overlapping rs182089148 that contacts the promoters of *LZTFL1* and *CCR1*. We observed that rs182089148-T led to opposite effects on the expression of *CCR1* (increase) and *CCR2* (decrease). The meta-analysis of the MPRA shows that the variant is MPRA active in CHRF cells. **C)** Locus plot for the variants impacting the genes *CCR7*, *SMARCE1* and *KRT10*. There are two independent signals: rs2228015, a missense and start lost variant in *CCR7*, and the index variant rs112401631, which sits in accessible chromatin in the lymphocyte and granulocyte monocyte lineages and has a PCHi-C connection with the promoter of *CCR7* only in the lymphocyte cell lineage. The index variant rs112401631 has three DE genes including a significant upregulation of *CCR7*. Both variants decrease LYMPH# in carriers of the alternative allele. The meta-analysis of the MPRA shows that the variant is MPRA active in K-562, HL-60 and THP-1 cells. **D)** Locus plot for the variants impacting the gene *RASAL3* and the GWAS trait LYMPH#. Two of the variants (rs56209154 and rs142945276, Tier 2, CADD values 23.5-29.3) are coding variants affecting the *RASAL3* gene. The ATU analysis revealed that rs569022307-C entails an ATU event that results in a higher contribution of two retained intron transcripts to the pool of transcripts of *RASAL3*. The meta-analysis of the MPRA shows that the variant is not MPRA active in any cell line. **E)** Locus plot for the variant rs17758695 with ATU in the candidate gene *BCL2*. rs17758695 is a very pleiotropic variant sitting in accessible chromatin in the early progenitor and stem cell stages of haematopoiesis described before^5^. ATAC-seq shows accessible chromatin in progenitor cell types and the PCHi-C track for naive CD4+ T cells indicates that the variant overlaps a region that contacts the promoter of *BCL2*. We detect the novel observation that the variant led to ATU in the *BCL2* gene in neutrophils and naive T-CD4 cells. The meta-analysis of the MPRA shows that the variant is MPRA active in K-562 and CHRF cells. **F)** Locus plot for the variants impacting the gene *ANK1* and the GWAS trait RET#. Three of the variants (rs780729970, rs34664882 and rs142626656, respectively Tier 1, Tier 1 and Tier 2, CADD values 15-29.3) are coding variants affecting the *ANK1* gene whereas a fourth independent signal (rs113630867, Tier 1) is an e- and s-QTL in whole blood upregulating the expression of the gene. The ATU analysis reveals that rs149489081-G entails an ATU event that results in a lower contribution of a processed transcript to the pool of transcripts of *ANK1*. We hypothesise that this change leads to a gain-of-function at the gene level resulting in an increase in RET# in keeping with the fact that the common eQTL upregulates the expression of *ANK1* and increases RET#. The meta-analysis of the MPRA shows that the variant is MPRA active in K-562, CHRF, HL-60 and THP-1 cells. **G)** Locus plot for the variant rs150649461 (chr1:92925654 G > C). The conditionally independent variant C01P092925706 (chr1:92925706 ATTAGAG > A) overlapped some of the rs150649461 tiles in the MPRA. For these tiles we assayed all the single alternative allele and double alternative allele versions and found that all possible combinations were MPRA positive (see **Table S4**). The ‘Index’ variant rs150649461-C allele led to a significant upregulation of *GFI1* and *EVI5* and an ATU event that resulted in a lower contribution of a processed transcript to the pool of transcripts of *EVI5*. **H)** Locus plot for the variant rs35592432 impacting the genes *FOXP1* and *FOXP1*-*IT1* and the GWAS trait LYMPH#. ATAC-seq shows accessible chromatin in the lymphoid lineage and the PCHi-C tracks for naive CD4+ T and naïve B cells indicate that the variant overlaps a region that contacts the promoter of *FOXP1*. In naïve T-CD4 cells we observe that the allele rs35592432-C significantly increases the expression of the gene *FOXP1*-*IT1* and leads to an ATU event in the FOXP1 gene with lower contribution of a processed transcript to the pool of transcripts of the gene. The meta-analysis of the MPRA shows that the variant is MPRA active in K-562 cells. **I)** Locus plot for the variant rs150221884 which tags a coding proxy, rs138903557 (CADD value 24.7, Tier 2), in the candidate gene *MFSD2B*. The ATU analysis revealed that rs150221884-T entails an ATU event that results in a higher contribution of a nonsense mediated transcript to the pool of transcripts of *MFSD2B*. The meta-analysis of the MPRA shows that the variant is not MPRA active in any cell line. **J)** Locus plot for the variant rs543594419 which tags the regulatory proxy rs74624637 (Tier 3) for the gene *TYMP*. The ATU analysis revealed that the haplotypes rs543594419-T-rs74624637-T and rs543594419-C-rs74624637-T entail an ATU event that results in a lower contribution of a retained intron transcript to the pool of transcripts of *TYMP*. The meta-analysis of the MPRA shows that the variant is MPRA active in K-562 and HL-60 cells. **K)** Locus plot for the variant rs572476245 for which manual curation indicates that a different tissue was needed to ascertain its effect on the *THPO* gene (Thrombopoietin) since this hormone is produced in the liver and the kidney and not expressed by blood cell types. Two of the variants (rs78565404 and rs6141, both Tier 1, CADD values 3.19 and 2.42) are coding variants affecting the *THPO* gene. The meta-analysis of the MPRA shows that the variant is not MPRA active in any cell line. **L)** MPRA results for the variant rs139141690. The meta-analysis of the MPRA shows that the variant is MPRA active in K-562 and HL-60 cells. *See Table S2 for the breakdown of variant classification in tiers and Table S3 for the abbreviations of blood phenotypes. C.I. = Conditionally Independent, DE = Differential Gene Expression, ATU = Differential Transcript Usage, UTR3 = variant in the 3’ UTR region. LD = linkage disequilibrium, PCHi-C = Promoter Capture HiC. * p value <= 0.05, ** p value <= 0.01, *** p value <= 0.001, ATU additive log ratio model. The dashed line in the MPRA plots indicates the position of the variant within the sequence assayed.*

**Supplementary Figure 5. Cascading impact of rs35592432-C in the transcripts usage of the TF *FOXP1* and the expression of its target genes. A)** GenIE results for the knock-in (highlighted in green) and UDPs (rest of profiles) of rs35592432 in Kolf2 iPSCs. Deletions encompassing the variant significantly increase the abundance of FOXP1 mRNA. **B)** Successful editing of H1 hESCs with the rs35592432 - C allele led to the isolation of three reference (G/G) and three homozygous alternative (C/C) clone lines. The cells were then differentiated to monocytes and bulk RNA-seq was performed at the endpoint. **C)** ATU analysis on *FOXP1* revealed a tendency to an increase in the relative contribution of a transcript without a CDS (ENST00000615603) in the C/C clone lines, (DRIMSeq model, p-value=0.198). **D)** Heatmap of the 112 DE genes between the G/G and C/C genotypes. The clone lines cluster by their corresponding genotype. Overrepresentation analysis in DE genes showed a 41-fold higher proportion of FOXP1 targets than expected by random chance (ORA test, p-value 3.27 10^-07).

**Supplementary Figure 6 related to Figure 4 and Figure 5. rs139141690-A downregulates the hematopoietic TF CUX1 leading to gene expression changes in the hematopoietic stem cell homeostasis and megakaryocyte differentiation that are recapitulated *‘in vitro’*.** **A)** Prediction of TF motifs in the sequence stretch spanned by the 80 bp deletion for the rs139141690-G and rs139141690-A alleles. The dashed line represents the genomic coordinate for rs139141690. The point mutation rs139141690 > A elicits a change of PU.1 TF motif to a FOXM1 motif, while CHIP-seq in whole blood shows the occupancy of the motif by both TFs. Moreover, a gene set enrichment analysis (GSEA) in the INTERVAL whole blood carriers revealed a significant enrichment of the *HSC homeostasis* and *Blood coagulation* pathways in heterozygous carriers of the variant (p-value 0.042 and 0.025 respectively). **B)** Position weight matrices for the rs139141690-G and rs139141690-A alleles. **C)** Differential analysis for chromVAR deviation scores in clusters 1 and 3 for peaks carrying *CUX1* motifs and peaks carrying *RUNX1* motifs across the different genotypes. **D)** Overrepresentation of DE genes in the GRNs of *CUX1* and of *RUNX1*. **E)** DE genes of the GRN *RUNX1*, GRN *CUX1* and genes belonging to both GRNs. *TF = Transcription Factor, GRN = Gene Regulatory Network. See Table S3 for all the cell types and GWAS traits abbreviations and Table S2 for the breakdown of variant classification in tiers.*

**Table S1 related to Figure 1, 3, 4, 5 and Table1. Table of blood phenotypes, cell types and datasets used in this study.** We indicate the link between blood phenotypes and lineages as well as which cell types are relevant for which phenotypes and lineages.

**Table S2 related to Figure 1. Variants classified in the different prioritisation tiers.** All the variants are in GRCh37.

**Table S3 related to Figure 2. MPRA results per tile and cell type prior to the meta-analysis.** All the genomic coordinates are in GRCh37.

**Table S4 related to Figure 2. MPRA results per cell type after the meta-analysis across tiles.** All the genomic coordinates are in GRCh37.

**Table S5 related to Figure 2. Results of DE and ATU analysis in INTERVAL whole blood and BLUEPRINT immune cell isolates.**

**Table S6 related to Figure 1,2,3,4,5 and 6. Master table with all the variants and classifications and the per variant results of the MPRA and the RNA-seq analysis.** All the variants are in GRCh37.

**Table S7 related to Methods. Table of oligos and sgRNAs.**

4.	ENCODE K-562 RNA report [https://www.encodeproject.org/rnaget-report/?type=RNAExpression&file.assay_title=polyA+plus+RNA-seq&file.biosa\mple_ontology.classification=cell+line&file.biosample_ontology.term_name=K562&limit=all](https://www.encodeproject.org/rnaget-report/?type=RNAExpression&file.assay_title=polyA+plus+RNA-seq&file.biosa%5Cmple_ontology.classification=cell+line&file.biosample_ontology.term_name=K562&limit=all).

5.	Ulirsch, J.C., Lareau, C.A., Bao, E.L., Ludwig, L.S., Guo, M.H., Benner, C., Satpathy, A.T., Kartha, V.K., Salem, R.M., Hirschhorn, J.N., et al. (2019). Interrogation of human hematopoiesis at single-cell and single-variant resolution. Nat. Genet. *51*, 683–693.
